# Reduction of Short-Chain Fatty Acid-Producing Gut Microbiota Leads to Transition from Rapid-Eye-Movement Sleep Behavior Disorder to Parkinson’s Disease

**DOI:** 10.1101/2020.08.07.242453

**Authors:** Hiroshi Nishiwaki, Tomonari Hamaguchi, Mikako Ito, Tomohiro Ishida, Tetsuya Maeda, Kenichi Kashihara, Yoshio Tsuboi, Jun Ueyama, Teppei Shimamura, Hiroshi Mori, Ken Kurokawa, Masahisa Katsuno, Masaaki Hirayama, Kinji Ohno

## Abstract

Gut dysbiosis has been reported repeatedly in Parkinson’s disease (PD), but once in rapid-eye-movement sleep behavior disorder (RBD) from Germany. Abnormal aggregation of α-synuclein fibrils causing PD possibly starts from the intestine. RBD patients frequently develop PD. Early-stage gut dysbiosis that is causally associated with PD is thus expected to be observed in RBD. We analyzed gut microbiota in 26 RBD patients and 137 controls by 16S rRNA-seq. Our RBD dataset was meta-analyzed with the German RBD dataset, and was compared with gut microbiota in 223 PD patients. Unsupervised clustering of gut microbiota by LIGER, a topic model-based tool for single-cell RNA-seq analysis, revealed four enterotypes in controls, RBD, and PD. Short-chain fatty acid (SCFA)-producing bacteria were conserved in an enterotype observed in controls and RBD, whereas they were less in enterotypes observed in PD. Genus *Akkermansia* and family *Akkermansiaceae* were consistently increased in both RBD in two countries and PD in five countries. No short-chain fatty acid (SCFA)-producing bacteria were significantly changed in RBD in two counties. In contrast, we previously reported that recognized and putative SCFA-producing genera *Faecalibacterium*, *Roseburia*, and *Lachnospiraceae ND3007 group* were consistently decreased in PD in five countries. Increased mucin-layer-degrading genus *Akkermansia* possibly accounts for the development of RBD, and an additional decrease of SCFA-producing genera is likely to be associated with the transition from RBD to PD.

**Importance:** Nineteen studies have been reported on gut microbiota in PD, whereas only one study has been reported in RBD from Germany. RBD has the highest likelihood ratio to develop PD. Our meta-analysis of RBD in Japan and Germany revealed increased mucin-layer-degrading genus *Akkermansia* in RBD. Genus *Akkermansia* may increase the intestinal permeability, as we previously observed in PD patients, and make the intestinal neural plexus exposed to oxidative stress, which can lead to abnormal aggregation of prion-like α-synuclein fibrils in the intestine. In contrast to PD, SCFA-producing bacteria were not decreased in RBD. As SCFA induces Treg cells, a decrease of SCFA-producing bacteria may be a prerequisite for the development of PD. We propose that prebiotic and/or probiotic therapeutic strategies to increase the intestinal mucin layer and to increase intestinal SCFA potentially retard the development of RBD and PD.

## Introduction

Parkinson’s disease (PD) is a progressive neurodegenerative disease exhibiting four major motor deficits of tremor, slowness of movement, rigidity, and postural instability (1). PD also exhibits non-motor symptoms that are characterized by dysautonomia (constipation, vomiting, orthostatic hypotension, excessive sweating, and dysuria) and mental disorders (depression, anxiety disorder, visual hallucination, and dementia) (1). Turning our eyes to the pathophysiology of PD, PD is caused by loss of the dopaminergic neurons in the substantia nigra, which is induced by abnormally aggregated α-synuclein fibrils (Lewy bodies) in the neuronal cells. Abnormal aggregation of α-synuclein fibrils behaves like prions, and is propagated to other neuronal cells probably via synapses (2). Lewy bodies are also observed in the cerebral cortex, the lower brainstem (3), the olfactory bulb (4), the autonomic nervous system (5), the salivary glands (6), the skin (7), and the intestine (6, 8, 9). Abnormal aggregation of α-synuclein fibrils possibly initiates in the intestinal neural plexus and ascends to the substantia nigra (4). Constipation, rapid-eye-movement sleep behavior disorder (RBD), and depression are frequently predisposed to the development of motor symptoms in PD in this order, which is in accordance with the ascending α-synucleinopathy (1). A total of 19 studies had been reported by us (10, 11) and others (12–28) for gut microbiota in PD. Our recent report included the largest cohort of PD patients, and the development of a novel nonparametric meta-analysis method that was applied to analyze gut microbiota in PD in five countries (11). Our meta-analysis revealed that increased genus *Akkermansia* and decreased genera *Roseburia* and *Faecalibacterium* were shared in PD across countries. In addition, these taxonomic changes were independent of the confounding effects of constipation, body mass index (BMI), sex, age and catechol-O-methyl transferase (COMT) inhibitor intake.

RBD is characterized by dream-enactment behaviors during the rapid eye movement (REM) sleep, when normal people lose muscle tone, called a state of atonia (29). The prevalence of RBD is estimated to be 0.5 to 2 percent (30, 31). RBD frequently predisposes to α-synucleinopathy including PD, dementia with Lewy bodies (DLB), and multiple system atrophy (MSA) (29). RBD patients sometimes have subtle sensory, motor, and cognitive deficits, as well as constipation, before the onset of PD and other α-synucleinopathies (29). PD has been classified into three group according as the disease progresses: preclinical PD (no overt symptoms even in the presence of neurodegeneration), prodromal PD (overt symptoms but lacking the criteria of PD), and clinical PD (overt symptoms satisfying the criteria of PD) (32). RBD is the most dependable hallmark of prodromal PD (32). Similarly, the likelihood ratio of RBD to develop PD is as high as 130 (33). Thus, therapeutic intervention to prevent transition from RBD to PD has a potential to become a causative treatment for PD (34).

In contrast to as many as 19 studies reported on gut microbiota in PD as stated above, only one study has been reported on 21 RBD patients along with 76 PD patients from Germany (22). They reported that gut microbiota in RBD was similar to that in PD. We recently reported increased genus *Akkermansia*, and decreased short-chain fatty acid (SCFA)-producing taxa in PD in five countries including the German dataset (11, 22). We here performed 16S rRNA-seq analysis of 26 RBD patients and 137 controls. We also meta-analyzed our dataset with the German dataset using a nonparametric meta-analysis method that we developed previously to identify shared taxonomic changes between the two countries, and compared RBD-associated taxonomic changes in two countries with PD-associated changes in five countries.

## Results

### PCoA plot to analyze the overall composition of gut microbiota in controls, RBD, and PDs

We performed 16S rRNA-seq analysis of gut microbiota in 26 patients with idiopathic RBD and 137 healthy controls. All RBD patients were diagnosed by International Classification of Sleep Disorders Criteria-Third Edition (35). We conducted PCoA analysis of gut microbiota in controls and RBD in our dataset, as well as gut microbiota in our previously reported PD subjects (Hoehn & Yahr scales 1-5) (11). The centers of gravity moved from the upper left to the lower right with disease progression from controls, RBD, to Hoehn & Yahr scales 1-5 (Fig. 1A). RBD was positioned close to the mildest form of PD with Hoehn & Yahr scale 1.

**Figure 1.**
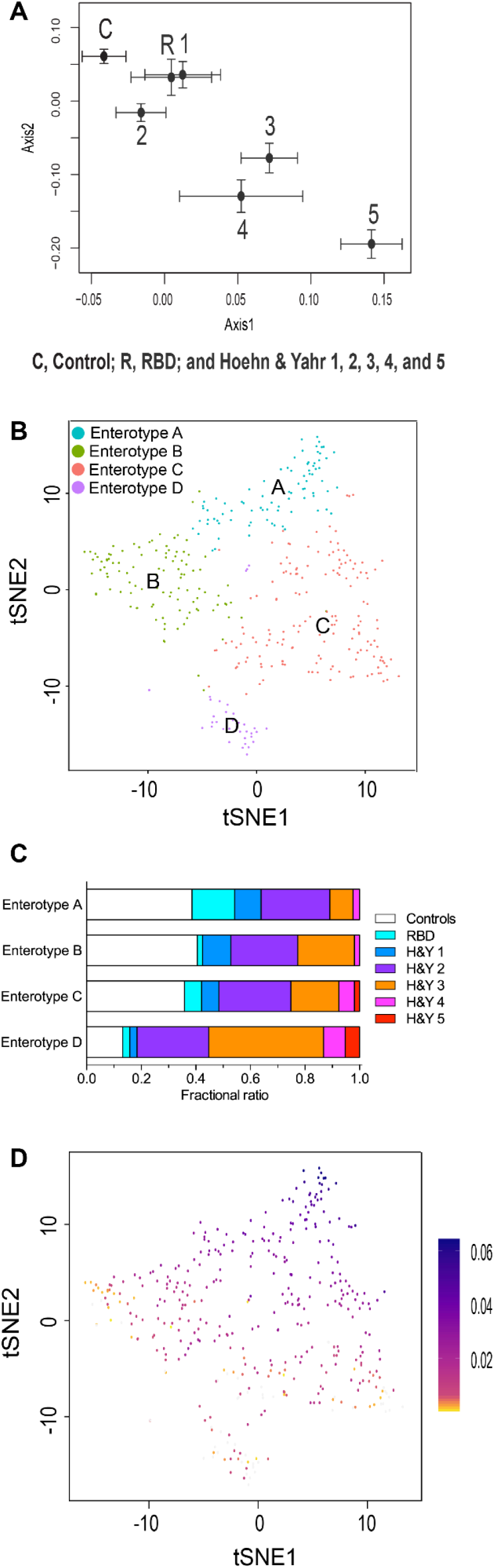
Overall compositions of gut microbiota in controls, RBD, and PD in our dataset. (A) PCoA plot showing the center of gravity of the overall compositions of gut microbiota in seven morbidity categories. The number of subjects in controls, RBD, Hoehn & Yahr scales 1-5 were 137, 26, 30, 99, 73, 16, and 5, respectively. Chao is used as a distance metric. Standard errors are indicated. (B) Unsupervised clustering of overall compositions of gut microbiota in controls, RBD, and PD by LIGER yielded four enterotypes. tSNE was adopted to visualize four clusters representing enterotypes A-D. (C) Fractional ratios of controls, RBD, and Hoehn & Yahr (H&Y) scales 1-5 in each enterotype. (D) Bacterial abundances in a total of 386 subjects were factorized into multiple factors. The first factor is color-coded in each subject on a tSNE plot indicated in (B). As SCFA-producing bacteria have high loadings in the first factor (see Table S1 in the supplemental material), individuals colored in blue carry a high proportion of SCFA-producing bacteria.

### LIGER analysis to reveal unsupervised enterotypes in a combined dataset of controls, RBD, and PD

We applied LIGER that was developed for topic model-based single-cell RNA-seq analysis (36) to make unsupervised clustering of gut microbiota in controls, RBD, and PD. Each cluster should represent an enterotype. LIGER revealed four enterotypes of A, B, C, and D (Fig. 1B). Examination of the proportion of controls, RBD, and Hoehn & Yahr scales 1-5 in each enterotype revealed that the proportion of controls was decreased in the order of enterotypes A to D, while the proportions of Hoehn & Yahr scales 3-5 were increased in the same order (Fig. 1C). The proportion of RBD was the highest in enterotype A. In factorization by LIGER, the fist factor contributes most to differentiate enterotypes A to D, and genera with high loadings in the first factor are major determinants of enterotypes. Color coding of the first factor in each subject showed a gradual decrease of the first factor from enterotypes A to D (Fig. 1D). The top ten genera with the highest loadings in the first factor are indicated in Table S1 in the supplemental material. It was interesting to note that, among the ten genera, nine produce SCFA and one putatively produces SCFA (*Lachnospiraceae ND3007 group*). Scatter plots of the ten genera in each enterotype showed that the abundances of these genera were also decreased in the order of enterotypes A to D (see Fig. S1 in the supplemental material). Among the ten genera, *Faecalibacterium*, *Roseburia*, and *Lachnospiraceae ND3007 group* were exactly the three genera that were decreased in PD in our previous meta-analysis of five countries (11). To summarize, unsupervised clustering of enterotypes revealed that enterotypes were shifted from A to D with transition from control, RBD, to Hoehn & Yahr scales 1-5, and that SCFA-producing genera were decreased from enterotypes A to D.

### Analysis of each taxon between controls and RBD in our dataset

We examined taxonomic differences between controls and RBD in our dataset using Analysis of Composition of Microbiomes (ANCOM) (37) and Wilcoxon rank sum test (Table S2a for the genus level analysis, and S2b for the family level analysis in the supplemental material). ANCOM was developed to reduce false discoveries by exploiting microbial compositional constraints (37, 38). The analyses revealed that seven genera were increased in RBD (*Ruminococcus 2*, *Alistipes*, *Akkermansia*, *Ruminococcaceae UCG-005*, *Ruminococcaceae UCG-004*, *[Eubacterium] coprostanoligenes group*, and *Family XIII AD3011 group*); two families were increased in RBD (*Rikenellaceae* and *Akkermansiaceae*); and no genera or families were decreased in RBD. The top ten increased or decreased genera in RBD by ANCOM were compared to those in PD in our previous report (11). Seven out of the top ten increased genera were shared between RBD and PD, whereas only one of the top ten decreased genera was shared between RBD and PD (Fig. 2).

**Figure 2.**
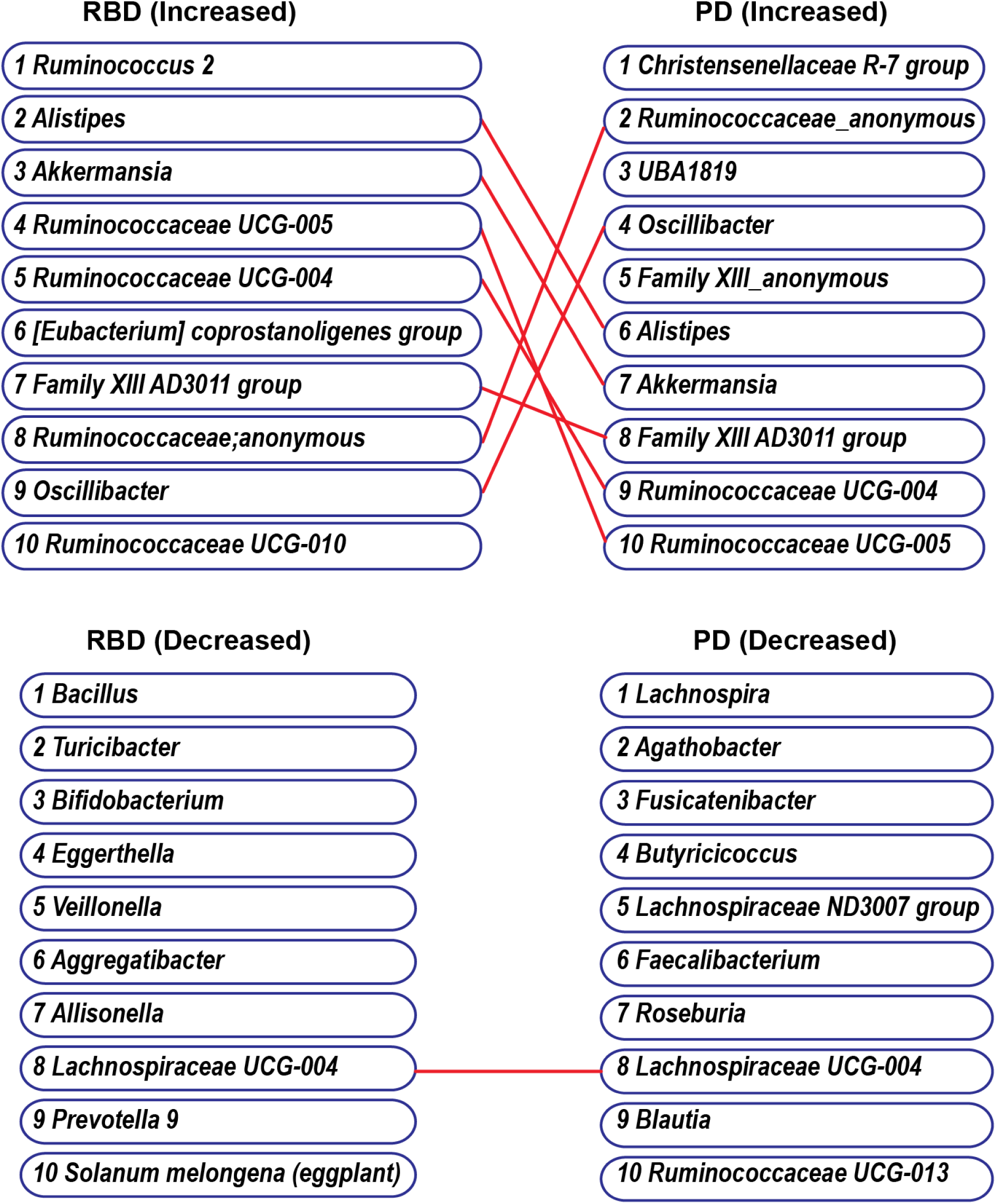
The 10 most increased and the 10 most decreased genera by ANCOM in RBD and PD both in our datasets. Identical genera are connected by a red line.

### Differences in demographic and clinical features between controls and RBD in our dataset

To search for possible confounding factors, we compared six features (age, sex, BMI, constipation, proton pump inhibitor (PPI), and H_2_ blocker) between RBD and controls in our dataset (Table 1). Compared to controls, RBD patients had higher ages and higher BMI, and included more males. Similarly, the ratios of constipation and PPI intake were higher in RBD patients.

**Table 1.**
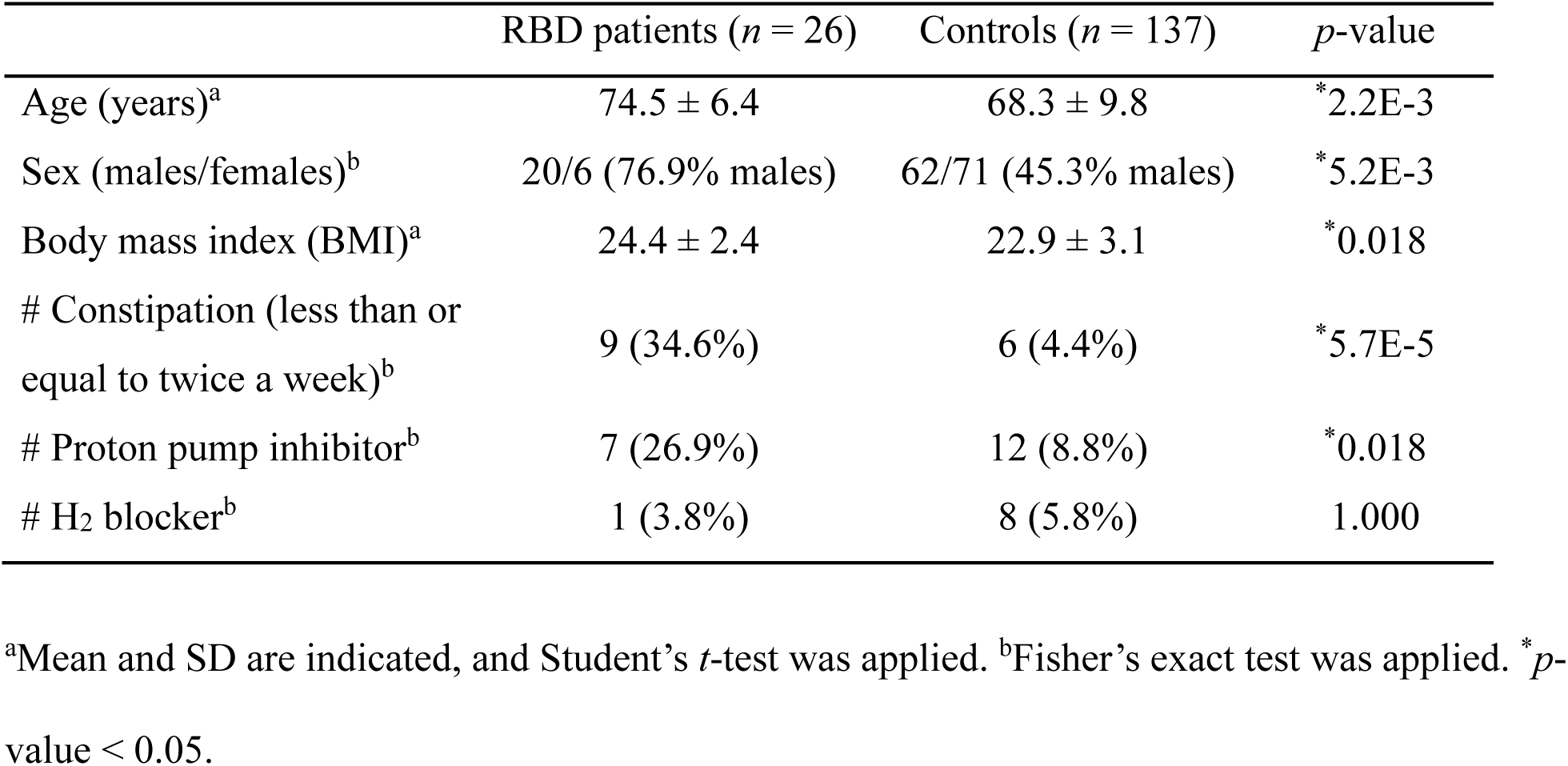
Demographic and clinical features of RBD and controls in our dataset

|  | RBD patients (*n* = 26) | Controls (*n* = 137) | *p*-value |
| --- | --- | --- | --- |
| Age (years)<sup>a</sup> | 74.5 ± 6.4 | 68.3 ± 9.8 | *2.2E-3 |
| Sex (males/females)<sup>b</sup> | 20/6 (76.9% males) | 62/71 (45.3% males) | *5.2E-3 |
| Body mass index (BMI)<sup>a</sup> | 24.4 ± 2.4 | 22.9 ± 3.1 | *0.018 |
| # Constipation (less than or equal to twice a week)<sup>b</sup> | 9 (34.6%) | 6 (4.4%) | *5.7E-5 |
| # Proton pump inhibitor<sup>b</sup> | 7 (26.9%) | 12 (8.8%) | *0.018 |
| # H<sub>2</sub> blocker<sup>b</sup> | 1 (3.8%) | 8 (5.8%) | 1.000 |

<sup>a</sup>Mean and SD are indicated, and Student's *t*-test was applied. <sup>b</sup>Fisher's exact test was applied. \**p*-value < 0.05.

### PERMANOVA to evaluate the differences in the overall composition of gut microbiota in controls and RBD, as well as in RBD and Hoehn & Yahr 1 scale

We next performed PERMANOVA to evaluate the difference in the overall composition of gut microbiota in controls and RBD (Table 2). The overall composition of gut microbiota between controls and RBD was statistically different by all three distance metrics, and the difference was not accounted for by age, sex, BMI, constipation, or PPI. Similarly, the overall composition of gut microbiota between RBD and Hoehn & Yahr scale 1 was also statistically different by Chao and weighted UniFrac but not by unweighted UniFrac. Again, the difference was not accounted for by any covariates.

**Table 2.**
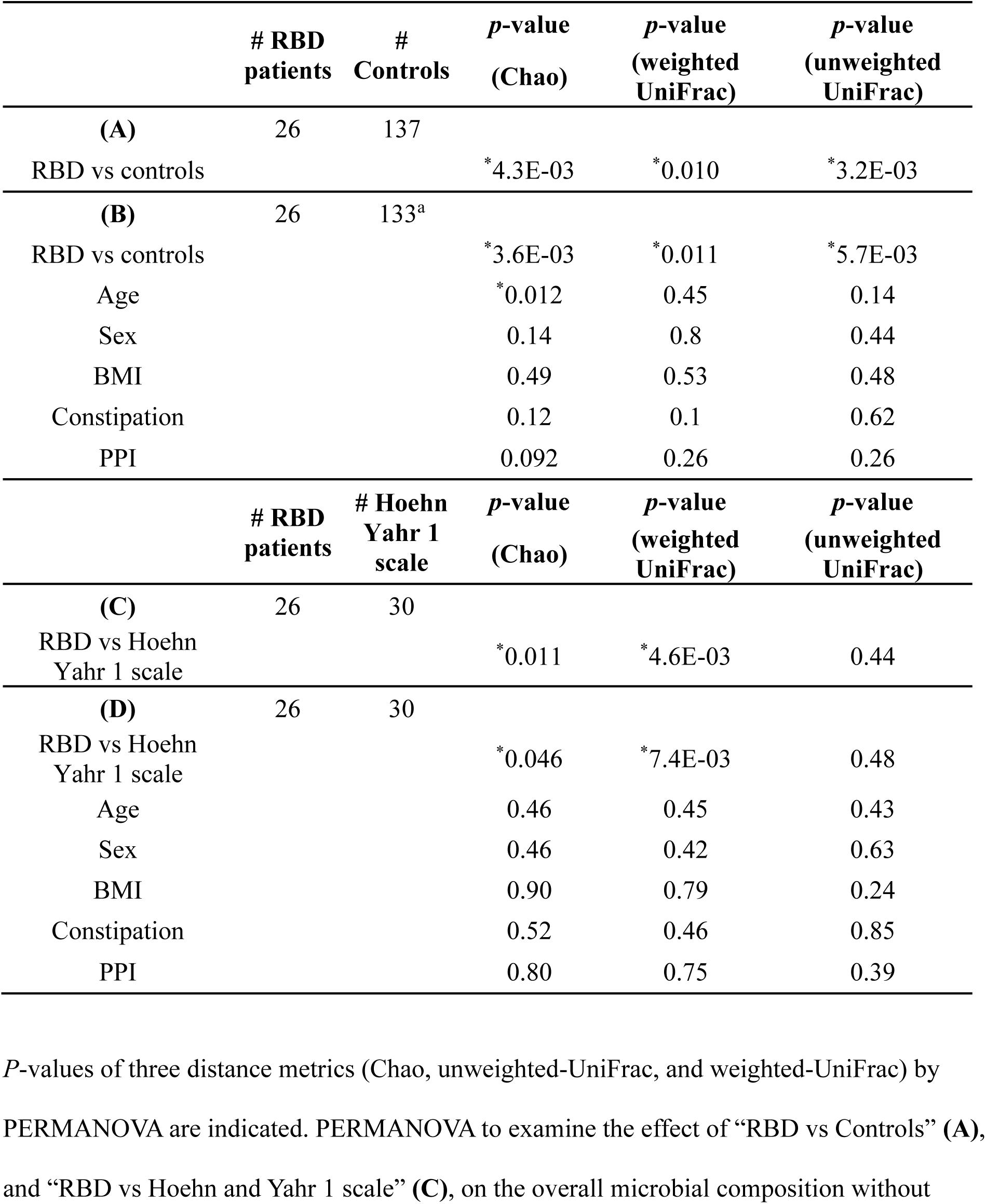
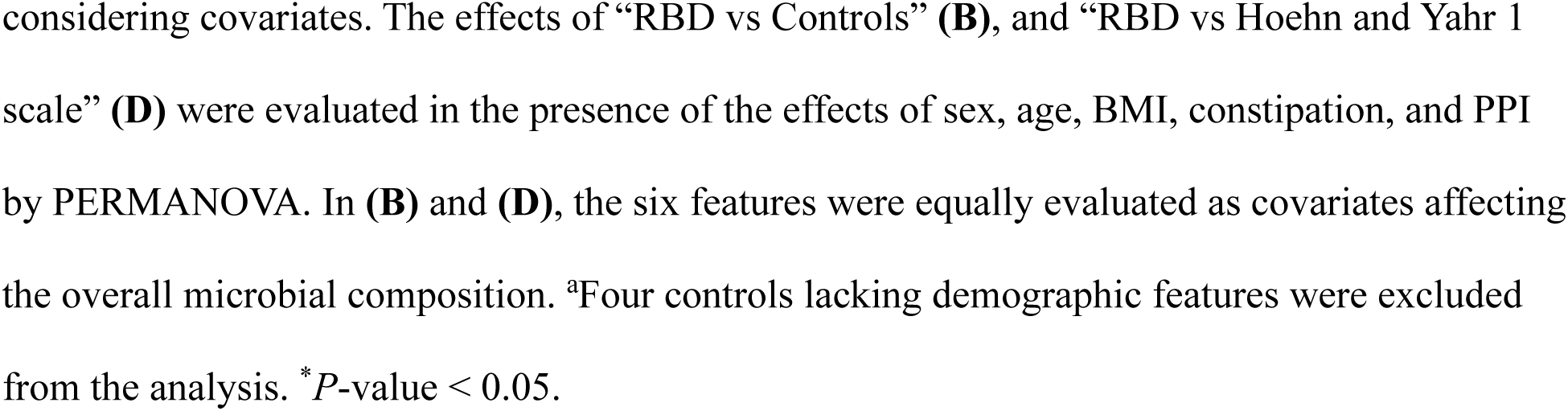
PERMANOVA to examine the effect of each factor on the overall bacterial composition in our dataset

|  | # RBD patients | # Controls | *p*-value (Chao) | *p*-value (weighted UniFrac) | *p*-value (unweighted UniFrac) |
| --- | --- | --- | --- | --- | --- |
| **(A)** | 26 | 137 |  |  |  |
| RBD vs controls |  |  | *4.3E-03 | *0.010 | *3.2E-03 |
| **(B)** | 26 | 133<sup>a</sup> |  |  |  |
| RBD vs controls |  |  | *3.6E-03 | *0.011 | *5.7E-03 |
| Age |  |  | *0.012 | 0.45 | 0.14 |
| Sex |  |  | 0.14 | 0.8 | 0.44 |
| BMI |  |  | 0.49 | 0.53 | 0.48 |
| Constipation |  |  | 0.12 | 0.1 | 0.62 |
| PPI |  |  | 0.092 | 0.26 | 0.26 |
|  | # RBD patients | # Hoehn Yahr 1 scale | *p*-value (Chao) | *p*-value (weighted UniFrac) | *p*-value (unweighted UniFrac) |
| **(C)** | 26 | 30 |  |  |  |
| RBD vs Hoehn Yahr 1 scale |  |  | *0.011 | *4.6E-03 | 0.44 |
| **(D)** | 26 | 30 |  |  |  |
| RBD vs Hoehn Yahr 1 scale |  |  | *0.046 | *7.4E-03 | 0.48 |
| Age |  |  | 0.46 | 0.45 | 0.43 |
| Sex |  |  | 0.46 | 0.42 | 0.63 |
| BMI |  |  | 0.90 | 0.79 | 0.24 |
| Constipation |  |  | 0.52 | 0.46 | 0.85 |
| PPI |  |  | 0.80 | 0.75 | 0.39 |

*P*-values of three distance metrics (Chao, unweighted-UniFrac, and weighted-UniFrac) by PERMANOVA are indicated. PERMANOVA to examine the effect of “RBD vs Controls” (A), and “RBD vs Hoehn and Yahr 1 scale” (C), on the overall microbial composition without
considering covariates. The effects of “RBD vs Controls” (**B**), and “RBD vs Hoehn and Yahr 1 scale” (**D**) were evaluated in the presence of the effects of sex, age, BMI, constipation, and PPI by PERMANOVA. In (**B**) and (**D**), the six features were equally evaluated as covariates affecting the overall microbial composition. <sup>a</sup>Four controls lacking demographic features were excluded from the analysis. \**P*-value < 0.05.

### Possible confounding factors in our dataset for nine taxa that were significantly changed in RBD in our dataset

We next asked whether any of the nine taxonomic changes in RBD were due to confounding factor(s). We thus performed GLMM analysis with constipation, BMI, sex, age, and PPI. We found that five genera (*Ruminococcus 2*, *Alistipes*, *Akkermansia*, *Ruminococcaceae UCG-004*, and *Family XIII AD3011 group*) and two families (*Rikenellaceae* and *Akkermansiaceae*) were changed in RBD after adjusting for constipation, BMI, sex, age, and PPI (Fig. 3 and bold letters in Table 3). In contrast, two genera (*Ruminococcaceae UCG-005* and *Family XIII AD3011 group*) were increased by age (Fig. 3 and underlines in Table 3). Three genera (*Akkermansia*, *Ruminococcaceae UCG-004* and *Family XIII AD3011 group*) and one family (*Akkermansiaceae*) were decreased by BMI (Fig. 3 and underlines in Table 3). Two genera (*Ruminococcaceae UCG-004* and *Family XIII AD3011 group*) were increased by constipation (Fig. 3 and underlines in Table 3).

**Figure 3.**
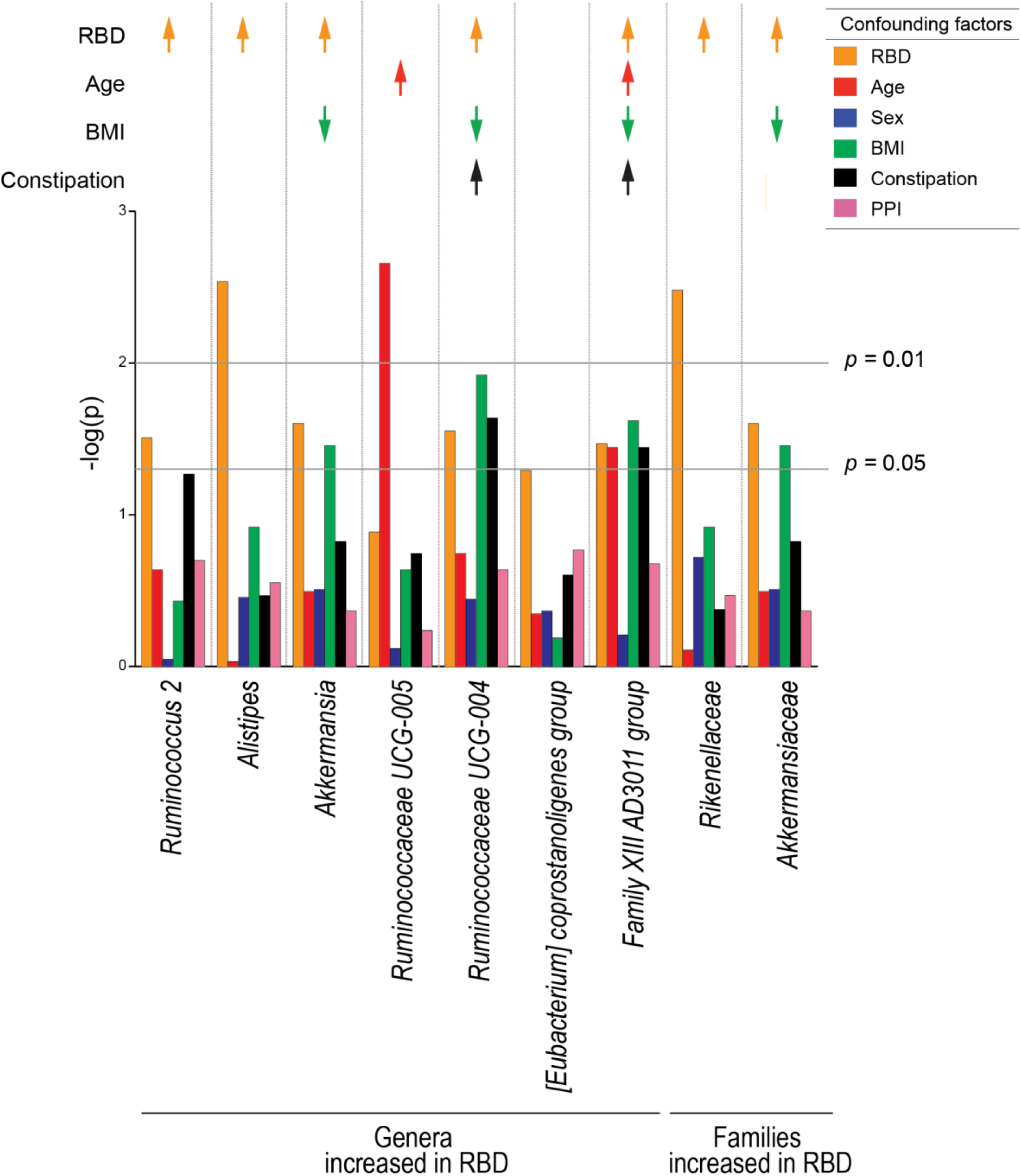
Generalized linear mixed model (GLMM) analysis to evaluate confounding factors of seven genera and two families that were significantly changed between RBD and controls in our dataset. The effects of RBD, age, sex, body mass index (BMI), constipation, and PPI were individually analyzed by mutually adjusting for covariates by GLMM. Arrows indicate taxa that were significantly changed by RBD (orange arrows), age (red arrows), BMI (green arrows) and constipation (black arrows) after adjusting for the other confounding factors. Upward and downward arrows indicate increased and decreased taxa, respectively. Exact *p*-values are indicated in Table 3.

**Table 3.**
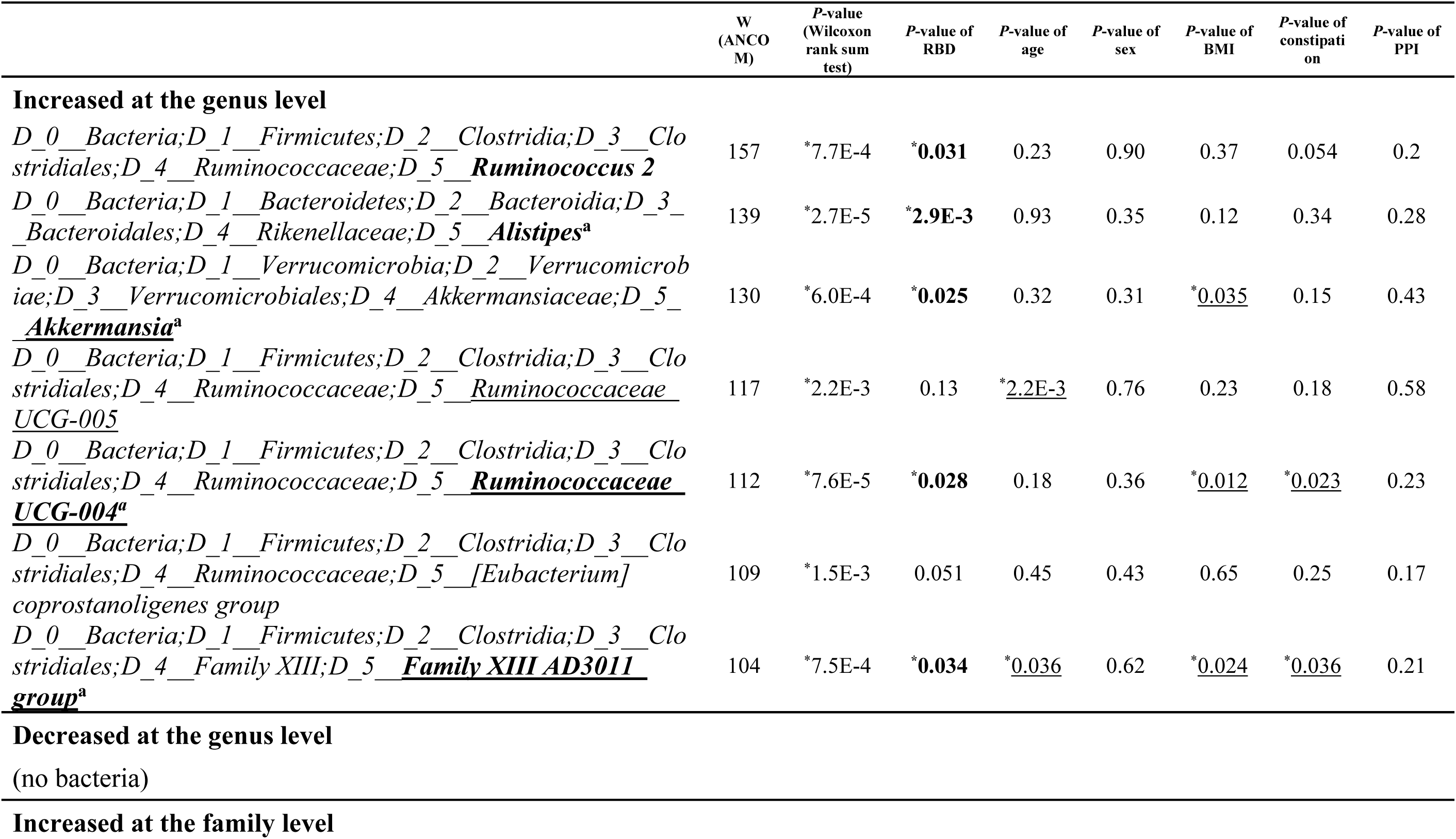
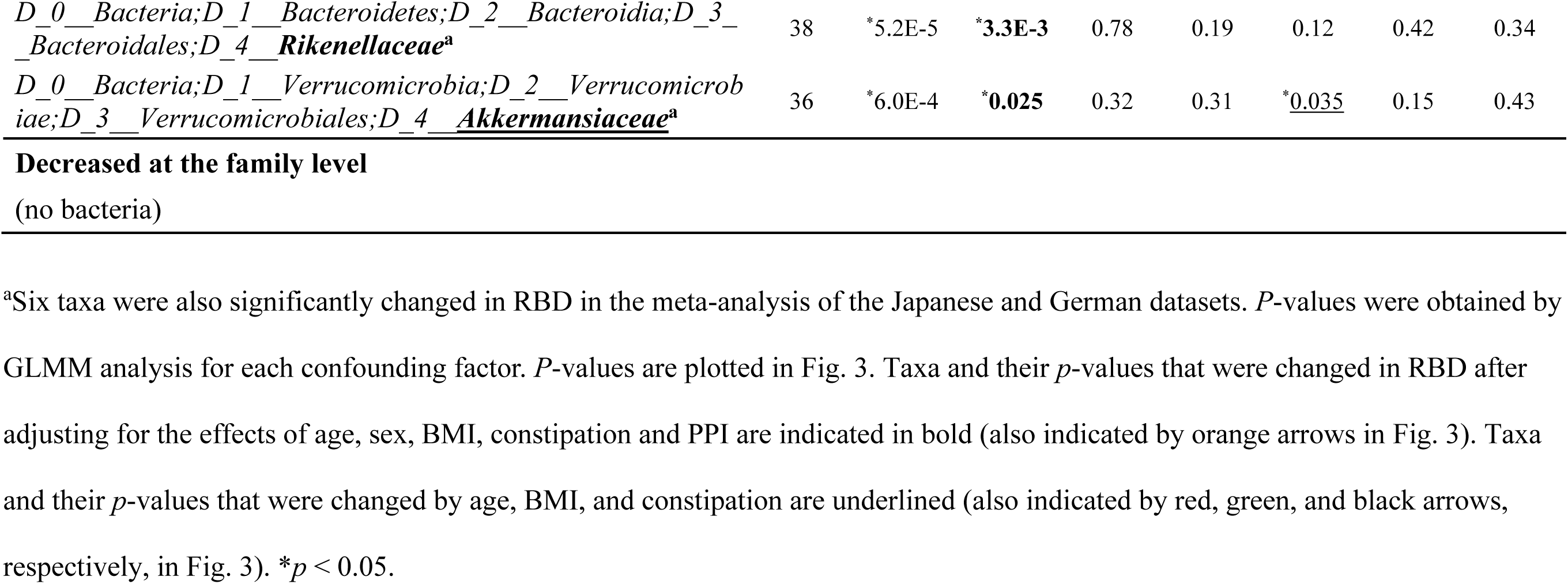
Generalized linear mixed model (GLMM) analysis of our dataset for nine bacterial taxa that were significantly changed in RBD in our dataset

### Meta-analysis of the Japanese and German datasets

Meta-analysis of gut microbiota was performed using the Japanese and German datasets (22). The effect size and relative abundance of 132 genera and 39 families are collated in Tables S3a and S3b, respectively, in the supplemental material. Our putative criteria (*I*^2^ < 25% and *p*-values of both FEM and REM after Bonferroni correction < 0.05) showed that four genera (*Ruminococcaceae UCG-004, Alistipes, Family XIII AD3011 group*, and *Akkermansia*) and two families (*Rikenellaceae* and *Akkermansiaceae*) were increased in RBD (Fig. 4, Table 4). These six taxa were a subset of the seven taxa that were significantly changed in RBD after adjusting for confounding factors in our dataset (Fig. 3 and bold letters in Table 3). Among the seven taxa, genus *Ruminococcus 2* was increased in Japan but not in Germany, and was excluded from forest plots (Fig. 4A). Forest plots of the six taxa in RBD in two countries along with in PD in five countries showed that all taxa tended to be increased in PD, and the most homogenous and significant increases were observed in genus *Akkermansia* and family *Akkermansiaceae* (Fig. 4A).

**Figure 4.**
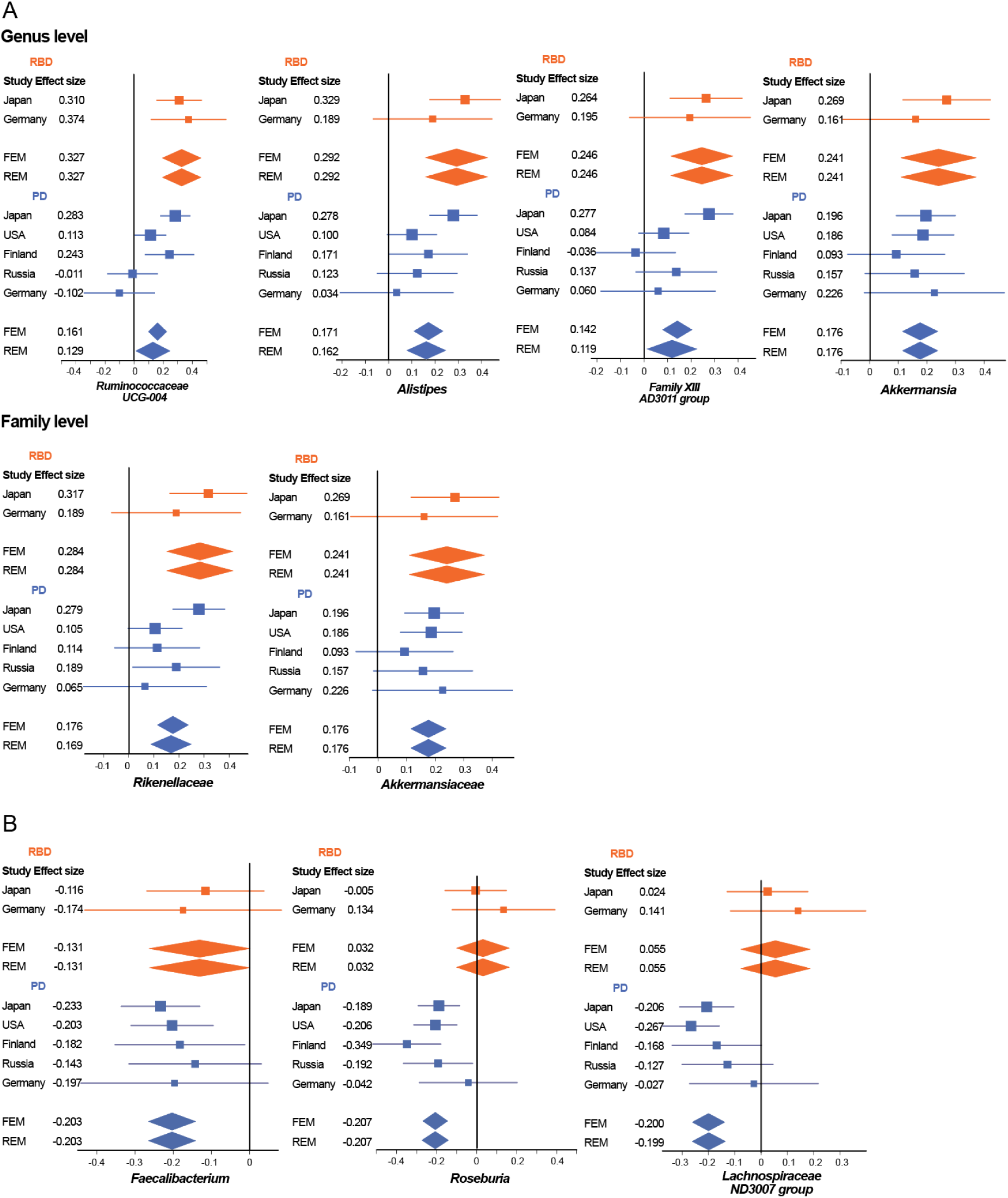
**(A)** Forest plots of four genera and two families that were significantly and homogenously changed in RBD in the Japanese and German datasets. Forest plots of PD in five datasets are also indicted in parallel. **(B)** Forest plots of two recognized and one putative SCFA-producing genera that were significantly and homogenously decreased in PD in five countries in our previous report (11). Forest plots of RBD in the Japanese and German datasets are also indicted in parallel. An effect size of each dataset, as well as the overall effect sizes by the fixed-effects model (FEM) and the random-effects model (REM), are indicated. Both lines and diamonds indicate 95% confidence intervals. Orange and blue symbols represent RBD and PD, respectively.

**Table 4.**
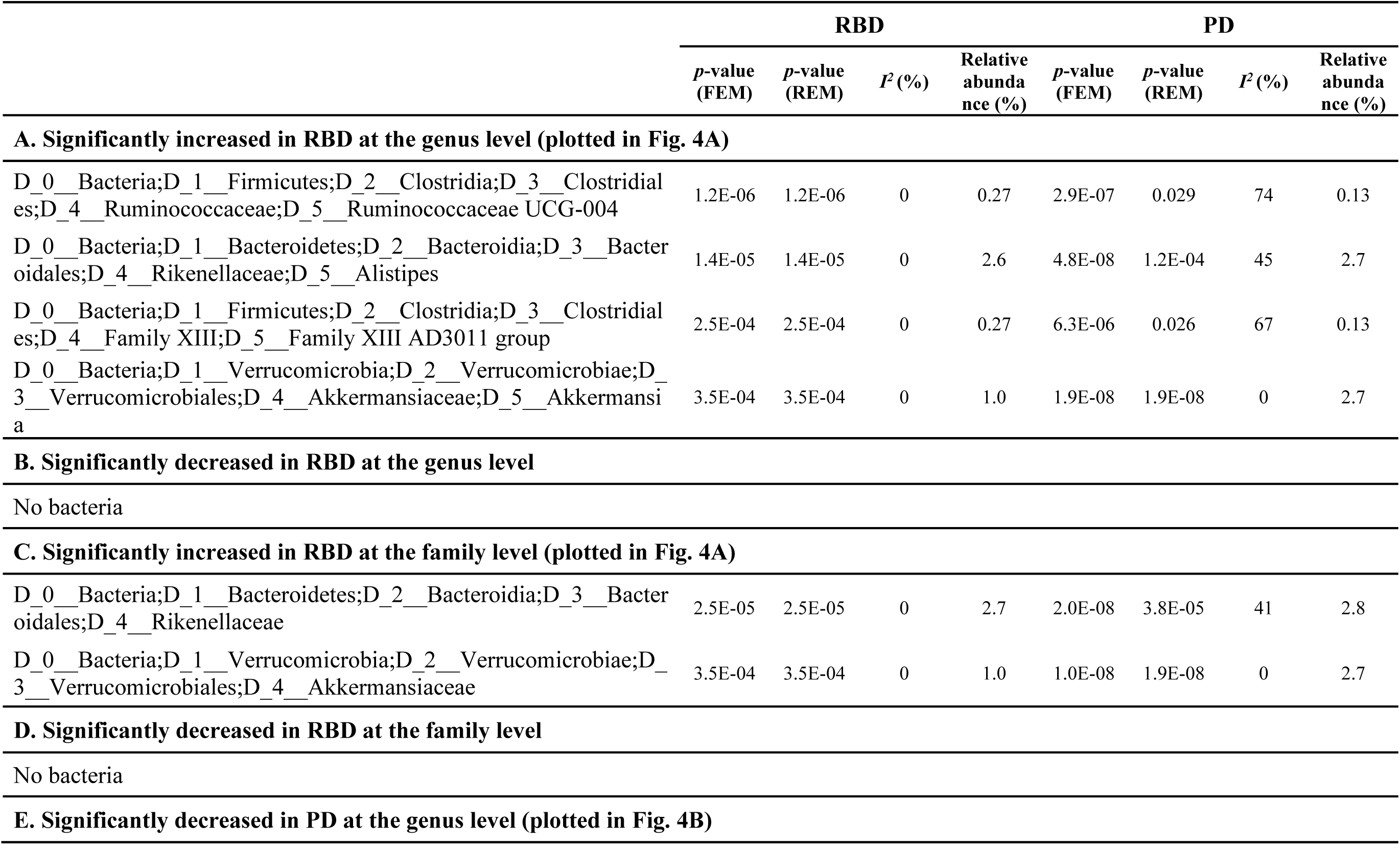
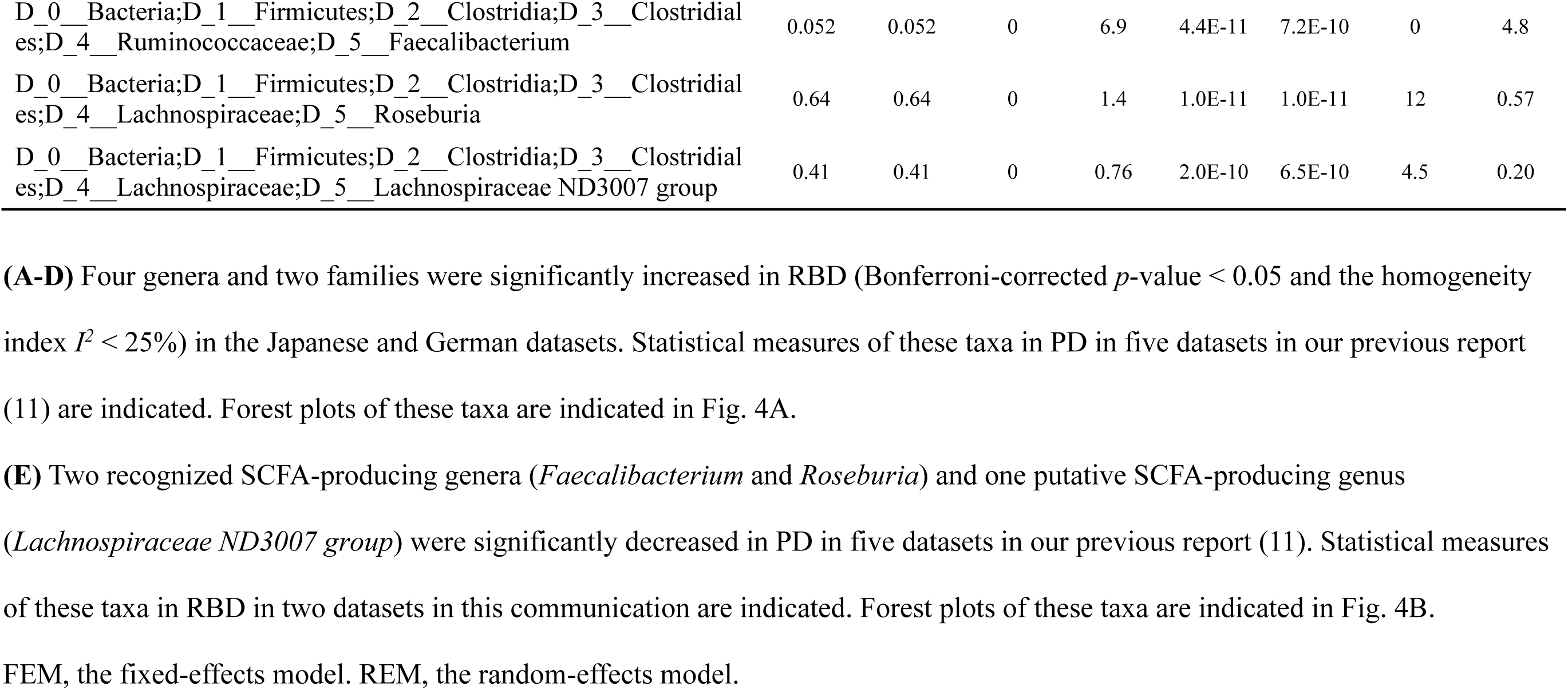
Statistical measures of meta-analysis of bacterial taxa in RBD in two datasets (Japan and Germany) and in PD in five datasets (Japan, USA, Finland, Russia, and Germany).

|  | RBD |  |  |  | PD |  |  |  |
| --- | --- | --- | --- | --- | --- | --- | --- | --- |
|  | *p*-value<br>(FEM) | *p*-value<br>(REM) | *I*<sup>2</sup> (%) | Relative<br>abundance (%) | *p*-value<br>(FEM) | *p*-value<br>(REM) | *I*<sup>2</sup> (%) | Relative<br>abundance (%) |
| **A. Significantly increased in RBD at the genus level (plotted in Fig. 4A)** |  |  |  |  |  |  |  |  |
| D_0__Bacteria;D_1__Firmicutes;D_2__Clostridia;D_3__Clostridiales;D_4__Ruminococcaceae;D_5__Ruminococcaceae UCG-004 | 1.2E-06 | 1.2E-06 | 0 | 0.27 | 2.9E-07 | 0.029 | 74 | 0.13 |
| D_0__Bacteria;D_1__Bacteroidetes;D_2__Bacteroidia;D_3__Bacteroidales;D_4__Rikenellaceae;D_5__Alistipes | 1.4E-05 | 1.4E-05 | 0 | 2.6 | 4.8E-08 | 1.2E-04 | 45 | 2.7 |
| D_0__Bacteria;D_1__Firmicutes;D_2__Clostridia;D_3__Clostridiales;D_4__Family XIII;D_5__Family XIII AD3011 group | 2.5E-04 | 2.5E-04 | 0 | 0.27 | 6.3E-06 | 0.026 | 67 | 0.13 |
| D_0__Bacteria;D_1__Verrucomicrobia;D_2__Verrucomicrobiae;D_3__Verrucomicrobiales;D_4__Akkermansiaceae;D_5__Akkermansia | 3.5E-04 | 3.5E-04 | 0 | 1.0 | 1.9E-08 | 1.9E-08 | 0 | 2.7 |
| **B. Significantly decreased in RBD at the genus level** |  |  |  |  |  |  |  |  |
| No bacteria |  |  |  |  |  |  |  |  |
| **C. Significantly increased in RBD at the family level (plotted in Fig. 4A)** |  |  |  |  |  |  |  |  |
| D_0__Bacteria;D_1__Bacteroidetes;D_2__Bacteroidia;D_3__Bacteroidales;D_4__Rikenellaceae | 2.5E-05 | 2.5E-05 | 0 | 2.7 | 2.0E-08 | 3.8E-05 | 41 | 2.8 |
| D_0__Bacteria;D_1__Verrucomicrobia;D_2__Verrucomicrobiae;D_3__Verrucomicrobiales;D_4__Akkermansiaceae | 3.5E-04 | 3.5E-04 | 0 | 1.0 | 1.0E-08 | 1.9E-08 | 0 | 2.7 |
| **D. Significantly decreased in RBD at the family level** |  |  |  |  |  |  |  |  |
| No bacteria |  |  |  |  |  |  |  |  |
| **E. Significantly decreased in PD at the genus level (plotted in Fig. 4B)** |  |  |  |  |  |  |  |  |
| D_0__Bacteria;D_1__Firmicutes;D_2__Clostridia;D_3__Clostridiales;D_4__Ruminococcaceae;D_5__Faecalibacterium | 0.052 | 0.052 | 0 | 6.9 | 4.4E-11 | 7.2E-10 | 0 | 4.8 |
| D_0__Bacteria;D_1__Firmicutes;D_2__Clostridia;D_3__Clostridiales;D_4__Lachnospiraceae;D_5__Roseburia | 0.64 | 0.64 | 0 | 1.4 | 1.0E-11 | 1.0E-11 | 12 | 0.57 |
| D_0__Bacteria;D_1__Firmicutes;D_2__Clostridia;D_3__Clostridiales;D_4__Lachnospiraceae;D_5__Lachnospiraceae ND3007 group | 0.41 | 0.41 | 0 | 0.76 | 2.0E-10 | 6.5E-10 | 4.5 | 0.20 |

FEM, the fixed-effects model. REM, the random-effects model.

We previously reported that two SCFA-producing genera (*Faecalibacterium* and *Roseburia*) and one putative SCFA-producing genus (*Lachnospiraceae ND3007 group*) were decreased in PD across countries (11). We assumed that genus *Lachnospiraceae ND3007 group* is a putative SCFA producer, because most genera in family *Lachnospiraceae* produce SCFA. None of the three recognized or putative SCFA-producing genera were decreased in RBD in our meta-analysis. However, forest plots of the three genera showed that genus *Faecalibacterium* tended to be decreased in RBD, but genera *Roseburia* and *Lachnospiraceae ND3007 group* were not (Fig. 4B).

### Relative abundances of four genera in progression of α-synucleinopathy

Plots of relative abundances of genera *Akkermansia*, *Faecalibacterium*, *Roseburia* and *Lachnospiraceae ND3007 group* in controls, RBD, and Hoehn & Yahr scales 1-5 showed that genus *Akkermansia* gradually increased, and genera *Faecalibacterium, Roseburia*, and *Lachnospiraceae ND3007 group* gradually decreased with progression of α-synucleinopathy (Fig. 5). Comparison of controls and RBD showed that genus *Akkermansia* was significantly increased in RBD. In contrast, genus *Faecalibacterium*, but not *Roseburia* or *Lachnospiraceae ND3007 grou*p, tended to be decreased in RBD (Fig. 5).

**Figure 5.**
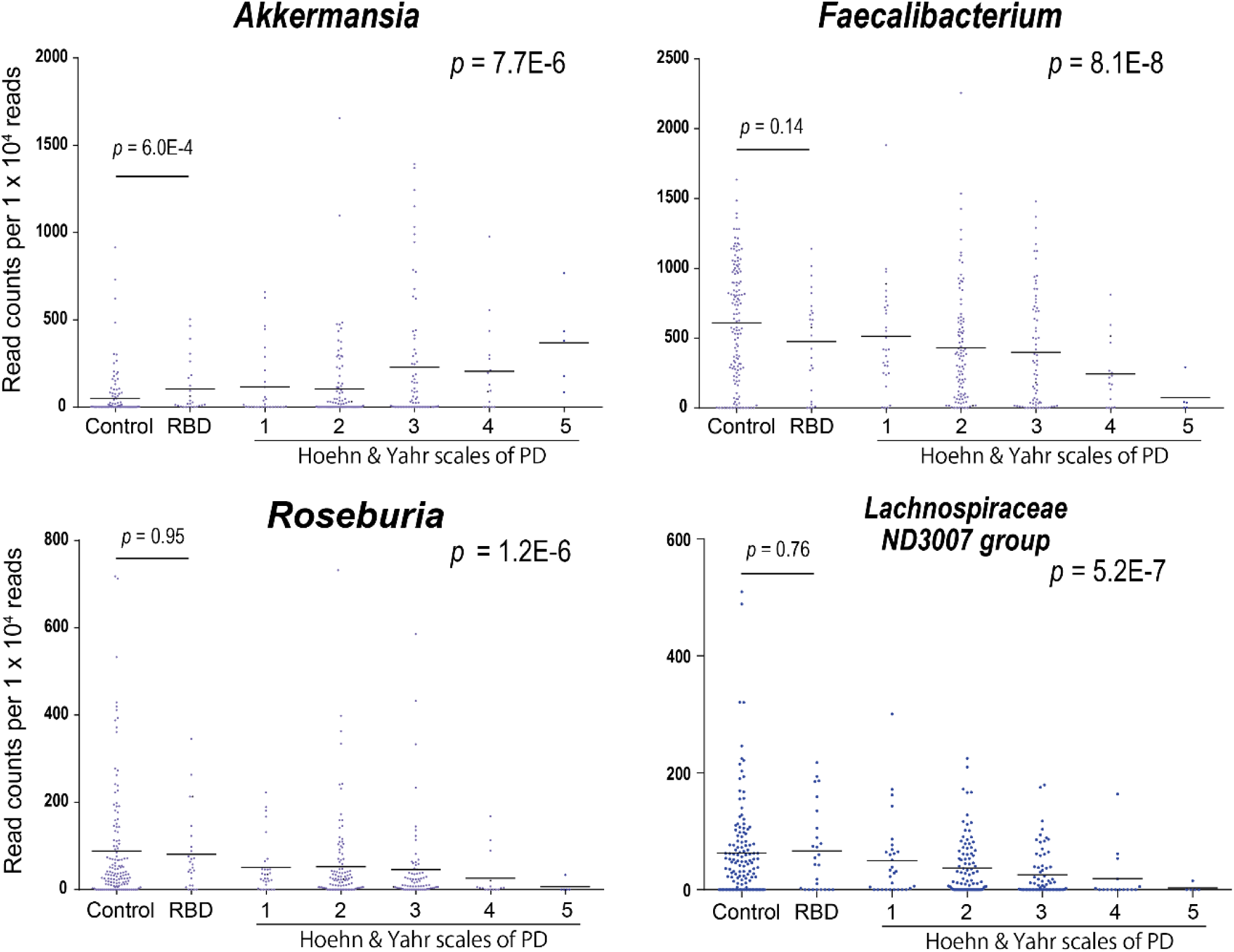
Reads counts of genera *Akkermansia*, *Faecalibacterium, Roseburia*, and *Lachnospiraceae ND3007 group* normalized for 1 x 10^4^ reads in controls, RBD, and Hoehn & Yahr scales1 to 5. Bar indicates an average in each category. *P*-values of Jonckheere-Terpstra trend test are shown on the right to indicate whether the genus increases or decreases monotonically. *P*-values of Wilcoxon rank sum test between controls and RBD (Table S2a in the supplemental material) are indicated on the left. Reads counts of genera *Akkermansia*, *Faecalibacterium, and Roseburia* in Hoehn & Yahr scales 1-5 were previously reported (11).

## Discussion

We analyzed gut microbiota in RBD in our dataset, and meta-analyzed the Japanese and Germany datasets (22). We first observed by PCoA analysis that the overall compositions of gut microbiota were gradually changing in controls, RBD, Hoehn & Yahr scales 1-5 in this order (Fig. 1A). We next applied LIGER (36) to 16S rRNA-seq for the first time. LIGER, which was developed for single-cell RNA-seq analysis, enables integrative non-negative matrix factorization by exploiting a topic model. Topic modeling that has been developed for text mining may be able to be applied to the analysis of gut microbiota (39, 40). LIGER revealed four enterotypes in controls, RBD, and Hoehn & Yahr scales 1-5 in an unsupervised manner (Fig. 1B). Enterotypes were shifted with transition from control, RBD, to Hoehn & Yahr scales 1-5 (Fig. 1C). SCFA-producing genera were similarly decreased with the shift in enterotypes (see Fig. S1 in the supplemental material). We showed that genus *Akkermansia* was increased in RBD in this communication and in PD in our previous report (11). Genus *Akkermansia*, however, was not detected in factorization by LIGER. Genus *Akkermansia* was likely to be underestimated by LIGER, because multiple SCFA-producing genera were coordinately decreased in PD, whereas genus *Akkermansia* was increased alone without any accompanying genera, which reduced a chance of detecting genus *Akkermansia* by topic modeling by LIGER. Both PCoA and LIGER indicate that gut dysbiosis advances with progression of α-synucleinopathy. Alternatively, patients with α-synucleinopathy with marked gut dysbiosis may progress faster than those with mild gut dysbiosis.

We additionally analyzed the overall compositions of gut microbiota in controls and RBD by PERMANOVA. Evaluation of the effects of covariates by PERMANOVA showed that the overall compositions of gut microbiota were statistically different between controls and RBD by all three distance metrics, and the difference was not due to covariates (Table 2). PERMANOVA similarly showed that the overall compositions of gut microbiota were statistically different between RBD and Hoehn & Yahr scale 1 by Chao and weighted UniFrac, but not by unweighted UniFrac (Table 2). Again, the difference was not due to covariates. Weighted UniFrac takes read counts into consideration to calculate the distance so that the effects of low-abundance taxa become small, whereas low-abundance taxa have more effects on unweighted UniFrac (41). Thus, RBD and Hoehn & Yahr scale 1 may have large differences in major taxa but not in minor taxa.

Analysis of individual taxa by ANCOM and the Wilcoxon rank sum test revealed that seven genera (see Tables S2a in the supplemental material) and two families (see Tables S2b in the supplemental material) were increased in RBD. Adjustment for possible confounding factors for the seven genera and two families by GLMM showed that increases of five genera (*Ruminococcus 2*, *Alistipes*, *Akkermansia*, *Ruminococcaceae UCG-004* and *Family XIII AD3011 group*) and two families (*Rikenellaceae* and *Akkermansiaceae*) were indeed accounted for by RBD (yellow arrows in Fig. 3), although age, BMI, and constipation had additional confounding effects on three genera (*Akkermansia*, *Ruminococcaceae UCG-004*, *Family XIII AD3011 group*) and one family (*Akkermansiaceae*). Among the five genera and two families, only genus *Akkermansia* and family *Akkermansiaceae* were also changed in PD in our meta-analysis of five countries (11).

Meta-analysis of the Japanese and German datasets revealed that four genera (*Ruminococcaceae UCG-004, Alistipes, Family XIII AD3011 group* and *Akkermansia*) and two families (*Rikenellaceae* and *Akkermansiaceae*) were increased in RBD (Fig. 4A, Table 4). Among these six taxa, we previously reported that genus *Akkermansia* and family *Akkermansiaceae* were consistently increased in PD across countries (11). We found that relative abundances of genus *Akkermansia* gradually increased from RBD to Hoehn & Yahr scales 1-5 (Fig. 5). *Akkermansia muciniphila* degrades the mucus layer of the gut (42), and erodes the mucus layer in the lack of dietary fibers (43). Indeed, intestinal permeability is increased in PD (44), and the serum lipopolysaccharide-binding protein levels are decreased in PD (10, 44). Reduced expression of a tight junction protein, occludin, in colonic biopsies in PD is similarly in accordance with the reduced mucus layer (45). Increased intestinal permeability may expose the intestinal neural plexus to oxidative stress and pesticide/herbicide (46). which subsequently allows the formation of abnormal α-synuclein aggregates in the intestine. Moreover, in the presence of other gut microbiota, *Akkermansia muciniphila* in mouse intestine enhances differentiation of follicular T cells, which mediate humoral immunity by B cells (47, 48). A high prevalence (20%) of autoimmune diseases in female patients with RBD compared to 5% in general population is in accordance with the *Akkermansia*-mediated increased humoral immunity (49). Similarly, RBD is sometimes associated with neuronal autoimmune diseases including narcolepsy, anti-IgLON5 disease, Kleine–Levin syndrome, multiple sclerosis, Guillain–Barré syndrome, anti-Ma2 encephalitis, LGI1 limbic encephalitis, Morvan’s syndrome, paraneoplastic cerebellar degeneration, and anti-NMDA receptor encephalitis (50).

Meta-analysis of the Japanese and German datasets also showed that no SCFA-producing genera were decreased in RBD (Fig. 4B, Table 4). We previously reported that three recognized and putative SCFA-producing genera (*Faecalibacterium*, *Roseburia*, and *Lachnospiraceae ND3007 group*) were consistently decreased in PD across countries (11). Although genus *Faecalibacterium* tended to be decreased in RBD, no significance was observed (Fig. 5). In contrast, genera *Roseburia* and *Lachnospiraceae ND3007 group* were not decreased in RBD (Fig. 5). Preservation of most of SCFA-producing bacteria in RBD was also implicated in the LIGER analysis, which showed that both controls and RBD were enriched in enterotype A (Fig. 1C), in which SCFA-producing bacteria were high (Fig. 1D). Major constituents of gut SCFAs, butyrate and propionate, induce anti-inflammatory Treg cells by inhibiting histone deacetylase (51, 52), and by binding to G protein-coupled receptors of GPR41, GPR43, and GPR109A (53, 54). Indeed, in mouse models of PD, SCFAs may (55–57) or may not (58) have beneficial effects on PD symptoms. In addition, in another German cohort, fecal SCFA concentrations were decreased in PD (14). Our analysis suggests that reduced fecal SCFA concentrations may be a prerequisite for the development of PD but not of RBD. Reduction of SCFA-producing bacteria culminating in the development of PD may start from genus *Faecalibacterium*. A decrease of genus *Faecalibacterium* may thus be a hallmark to predict transition from RBD to PD. We expect that administration of SCFA and probiotics/prebiotics to increase the intestinal SCFA possibly retard the progression of α-synucleinopathy at the stage of RBD.

## Materials and Methods

### Patients in our dataset

All studies were approved by the Ethical Review Committees of the Nagoya University Graduate School of Medicine (approval #2016-0151), Iwate Medical College Hospital (H28-123), Okayama Kyokuto Hospital (approval #kyoIR-2016002), and Fukuoka University, School of Medicine (approval #2016M027). We obtained written informed consent from all patients and controls.

We recruited 26 patients with idiopathic RBD and their 137 healthy controls from four hospitals to participate in this study from September 2015 to February 2018. Among the 137 healthy controls, 8 were healthy spouses of RBD patients. All RBD patients were diagnosed by International Classification of Sleep Disorders Criteria-Third Edition (35). Subjects with diabetes mellitus, heart failure, liver cirrhosis, any malignancy, hematological diseases, or autoimmune diseases were excluded from our study. Subjects who have taken any antibiotics in the past one month were similarly excluded.

### DNA isolation and 16S rRNA V3-V4 sequencing in our dataset

The detailed transportation procedures of a fecal sample from participant’s home to the Nagoya University, freeze-drying of the fecal sample (26), and DNA isolation were described previously (11). The V3–V4 hypervariable region of the bacterial 16S rRNA gene was amplified by primer 341F, 5’-CCTACGGGNGGCWGCAG-3’ and primer 805R, 5’-GACTACHVGGGTATCTAATCC-3’. Paired-end sequencing of 300-nucleotide fragments was performed using the MiSeq reagent kit V3 on a MiSeq System (Illumina). Taxonomic analysis was performed with QIIME2 (59). Operational taxonomic units (OTUs) were generated using DADA2 and the SILVA taxonomy database release 132 (60) was used for taxonomic identification.

### Analysis of the overall gut microbiota between controls, RBD, and PD in our dataset

For the overall analysis of gut microbiota, PD samples in our previous report (11) were included. We first performed principal coordinate analysis (PCoA) of each subject, and the centers of gravity and standard errors in seven categories of controls, RBD, and Hoehn & Yahr scales 1-5 were plotted.

We next employed the Linked Inference of Genomic Experimental Relationships (LIGER) (36), which uses integrative non-negative matrix factorization (iNMF) for single-cell RNA-seq analysis, for unsupervised clustering of gut microbiota of controls, RBD, and Hoehn & Yahr scales 1-5. LIGER enabled us to identify four enterotypes, each of which was comprised of a set of bacteria that were synchronously changed in each subject.

### Analysis of each taxon between controls and RBD in our dataset

Taxa were filtered at the genus and family levels using the following conditions. For each taxon, we counted the number of samples in which the relative abundance of the taxon of interest was greater than 1E-4. The number of such samples should be 17 or more (more than ∼10% of all samples). We thereby chose 50 families and 168 genera.

The difference in the abundance of each taxon between RBD and controls was analyzed by Analysis of Composition of Microbiomes (ANCOM) (37), as well as by the Wilcoxon rank-sum test. ANCOM was performed on R (https://github.com/antagomir/scripts/tree/master/R/ancom). The Wilcoxon rank-sum test was performed with the mannwhitneyu functionality of scipy.stat on Python 3.6.5. The threshold of W calculated in ANCOM was set to more than 0.6 × *N*, where *N* is the number of taxa. The difference in the abundance of each taxon between RBD and controls was also analyzed by the Wilcoxon rank-sum test followed by calculation of the false discovery rate (FDR) using the Benjamini-Hochberg procedure. The FDR threshold was set to 0.05. Bacterial taxa filtered for both W and FDR were assumed to be significant.

### Possible confounding factors in our dataset for nine taxa that were significantly changed in our dataset

Six demographic and clinical features [age, sex, body mass index (BMI), constipation, proton pump inhibitor intake, and H_2_ blocker intake] were compared between RBD and controls in our dataset using either Student’s *t*-test or Fisher’s exact test. Subjects with the stool frequency twice a week or less were defined to be constipated (61). Constipation and BMI were statistically different between RBD and controls.

We examined lack of multicollinearity between RBD, constipation, BMI, sex, and age by calculating the variance inflation factor (VIF) using the R package HH version 3.1-40. We verified that the VIFs were all less than 2, indicating that there was no multicollinearity between RBD, constipation, BMI, sex, and age.

Next, we analyzed the effects on the overall composition of gut microbiota of (i) RBD or controls, and (ii) RBD or controls, age, sex, BMI, constipation, and PPI in our dataset comprised of controls and RBD using PERMANOVA (62). We similarly analyzed the effects on the overall composition of gut microbiota of (iii) RBD or Hoehn & Yahr 1 scale, and (iv) RBD or Hoehn & Yahr 1 scale, age, sex, BMI, constipation, and PPI in a combined dataset comprised of our current RBD subjects and PD subjects with Hoehn & Yahr 1 scale in our previous report (11) using PERMANOVA (62). All genera were included in this analysis. The effects were evaluated by three distance metrics of Chao (63), unweighted-UniFrac (64), and weighted-UniFrac (64). Chao and unweighted/weighted-UniFrac distances were calculated using the R package vegan and QIIME2, respectively.

Seven genera and two families that were identified in our dataset were subjected to GLMM (Generalized Linear Mixed Model) analysis using the function “glmer.nb” of the R package lme4 by setting an option to accept taxonomic variations from subject to subject.

### Meta-analysis of the Japanese and German datasets

Our Japanese dataset was comprised of 26 RBD patients and 137 healthy controls, whereas the German dataset was comprised of 20 RBD patients and 38 healthy controls (22). We first collated the experimental methods and demographic features (see Table S4 in the supplemental material), as well as statistical measures of sequencing depths (see Table S5 in the supplemental material) of the two datasets. The read count of each sample was all more than 10,000 in the two datasets, and no sample was excluded from our meta-analysis. For each taxon, we counted the number of samples in which the relative abundance of the taxon was more than 1E-4. We then filtered 39 families and 132 genera, in which the number of such samples was more than 10% (17/163 and 6/58) in both datasets.

In the meta-analysis, we applied two criteria that we used in our previous report (11) to identify homogeneously and significantly changed taxa in the Japanese and German datasets. The two criteria were that *I*^2^, representing heterogeneity in meta-analysis, was below 25% (65) and that the *p*-values after Bonferroni correction for FEM and REM were both less than 0.05.

### Data availability

FASTQ files of our RBD dataset are available at the DNA Data Bank of Japan (DDBJ) under the accession number DRA009322 (https://www.ncbi.nlm.nih.gov/sra/?term=DRA009322). FASTQ files of our PD dataset was previously deposited at DDBJ under the accession number DRA009229 (https://www.ncbi.nlm.nih.gov/sra/?term=DRA009229).

## Acknowledgements

We thank Keiichi Takimoto, Anzu Suzuki, Rino Asai, Yukina Matsuzaki, Sayaka Inagaki, Yuka Mishima, Yurika Muramatsu, and Tomomi Yamada at the Nagoya University Graduate School of Medicine for preparing DNA from fecal samples.

---

**Supplementary Table S1.** The top 10 genera with the highest loadings in the first factor by LIGER analysis

| Genus | Factor loading |
| --- | --- |
| D_0__Bacteria;D_1__Firmicutes;D_2__Clostridia;D_3__Clostridiales;D_4__Lachnospiraceae;D_5__Fusicatenibacter | 33.0 |
| D_0__Bacteria;D_1__Firmicutes;D_2__Clostridia;D_3__Clostridiales;D_4__Lachnospiraceae;D_5__Lachnospiraceae UCG-004 | 29.6 |
| D_0__Bacteria;D_1__Firmicutes;D_2__Clostridia;D_3__Clostridiales;D_4__Ruminococcaceae;D_5__Faecalibacterium | 27.8 |
| D_0__Bacteria;D_1__Firmicutes;D_2__Clostridia;D_3__Clostridiales;D_4__Ruminococcaceae;D_5__Butyricicoccus | 26.4 |
| D_0__Bacteria;D_1__Firmicutes;D_2__Clostridia;D_3__Clostridiales;D_4__Lachnospiraceae;D_5__Anaerostipes | 26.2 |
| D_0__Bacteria;D_1__Firmicutes;D_2__Clostridia;D_3__Clostridiales;D_4__Lachnospiraceae;D_5__Lachnospiraceae ND3007 group | 24.9 |
| D_0__Bacteria;D_1__Bacteroidetes;D_2__Bacteroidia;D_3__Bacteroidales;D_4__Bacteroidaceae;D_5__Bacteroides | 24.7 |
| D_0__Bacteria;D_1__Firmicutes;D_2__Clostridia;D_3__Clostridiales;D_4__Lachnospiraceae;D_5__Roseburia | 24.6 |
| D_0__Bacteria;D_1__Firmicutes;D_2__Clostridia;D_3__Clostridiales;D_4__Lachnospiraceae;D_5__Blautia | 24.0 |
| D_0__Bacteria;D_1__Firmicutes;D_2__Clostridia;D_3__Clostridiales;D_4__Lachnospiraceae;D_5__Lachnospira | 23.3 |

**Supplementary Table S2a.**
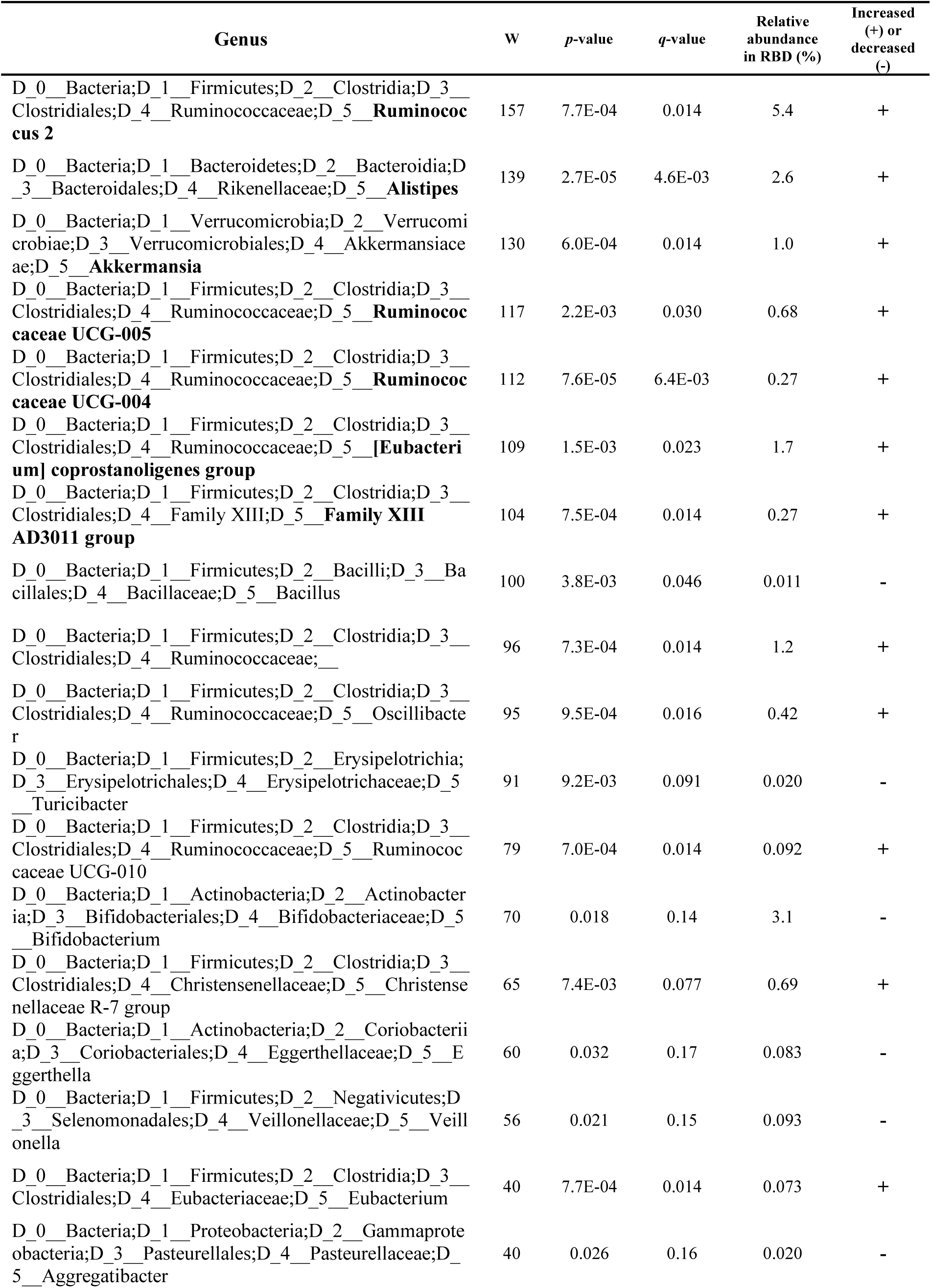
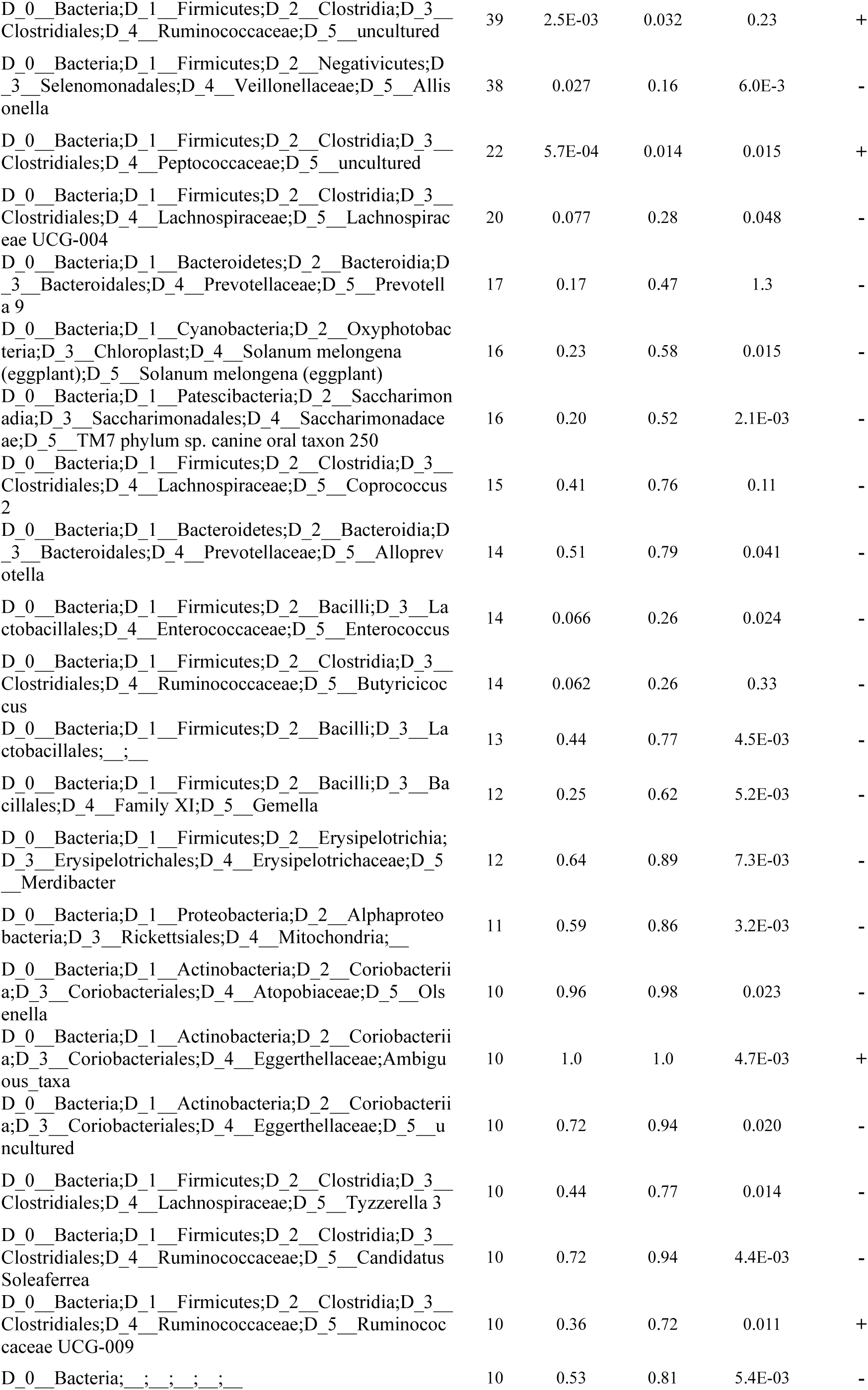
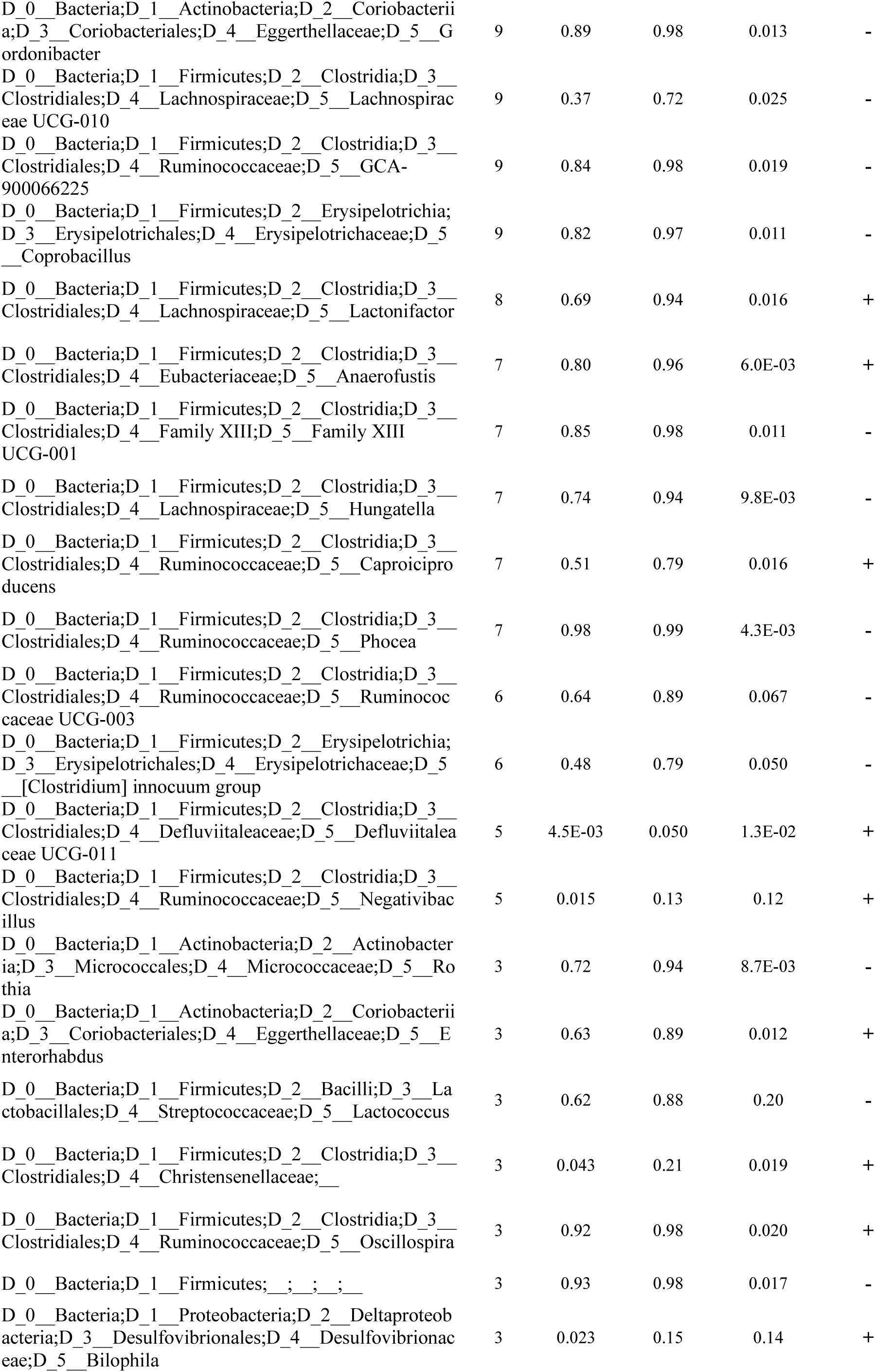
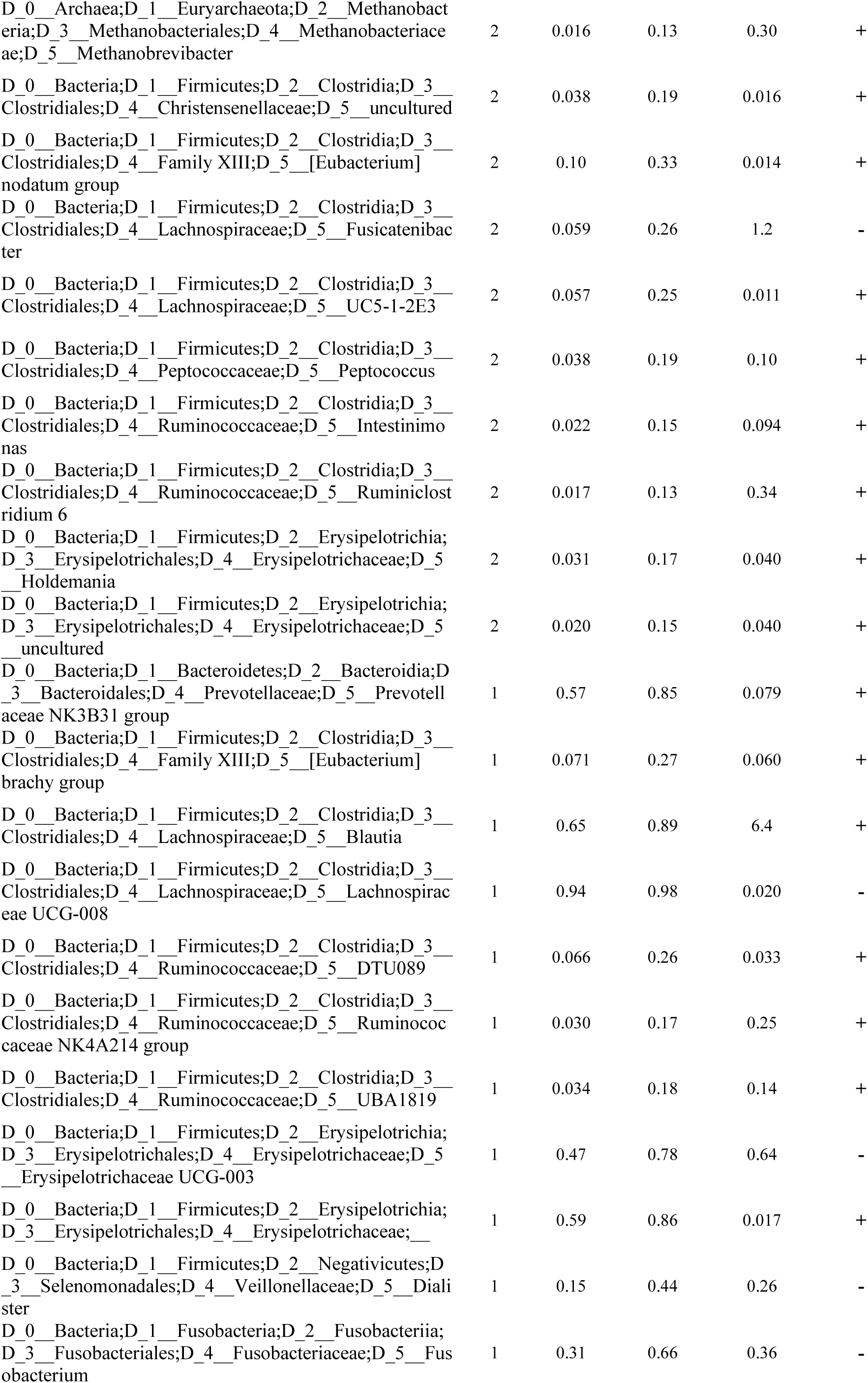
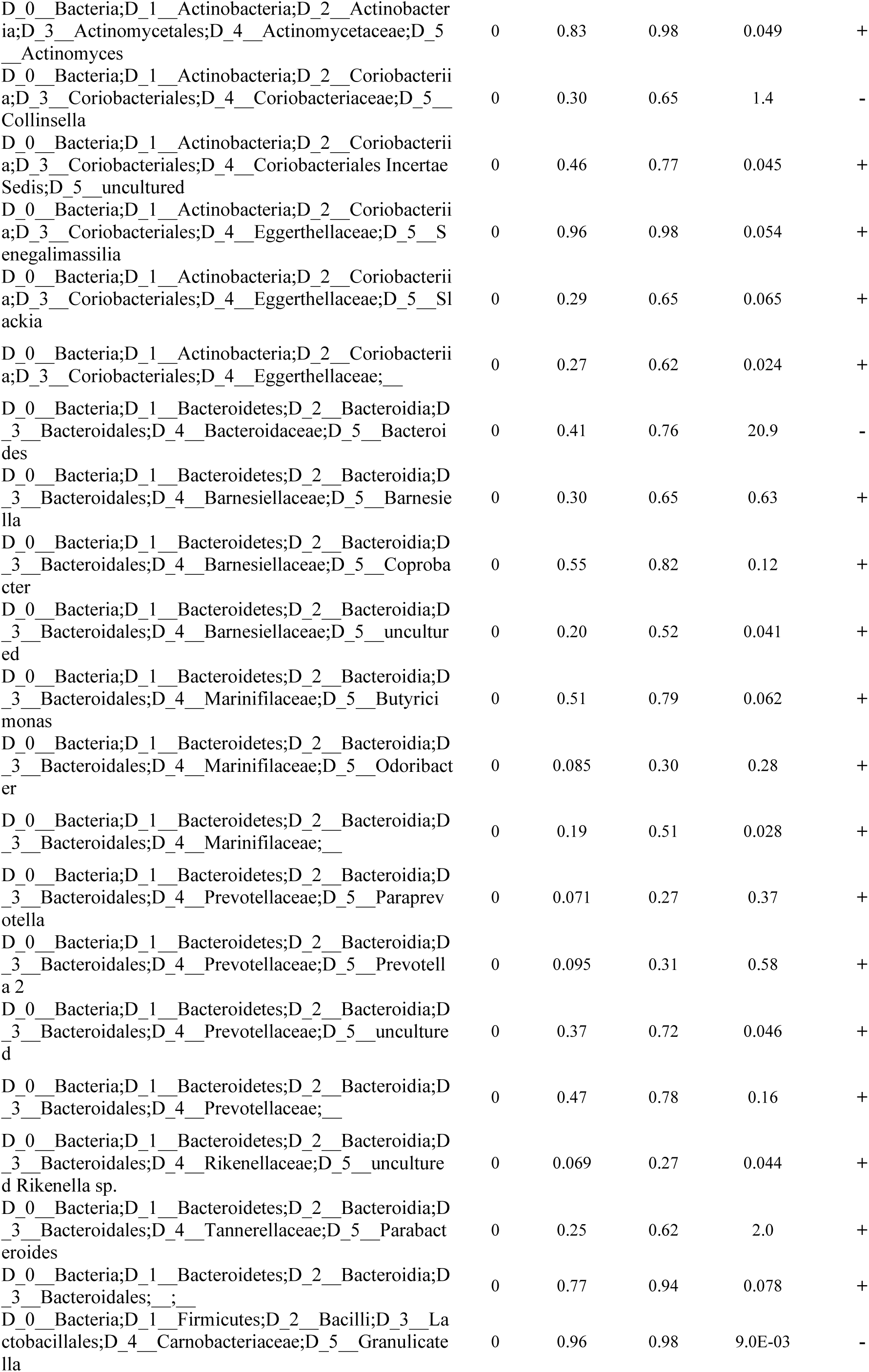
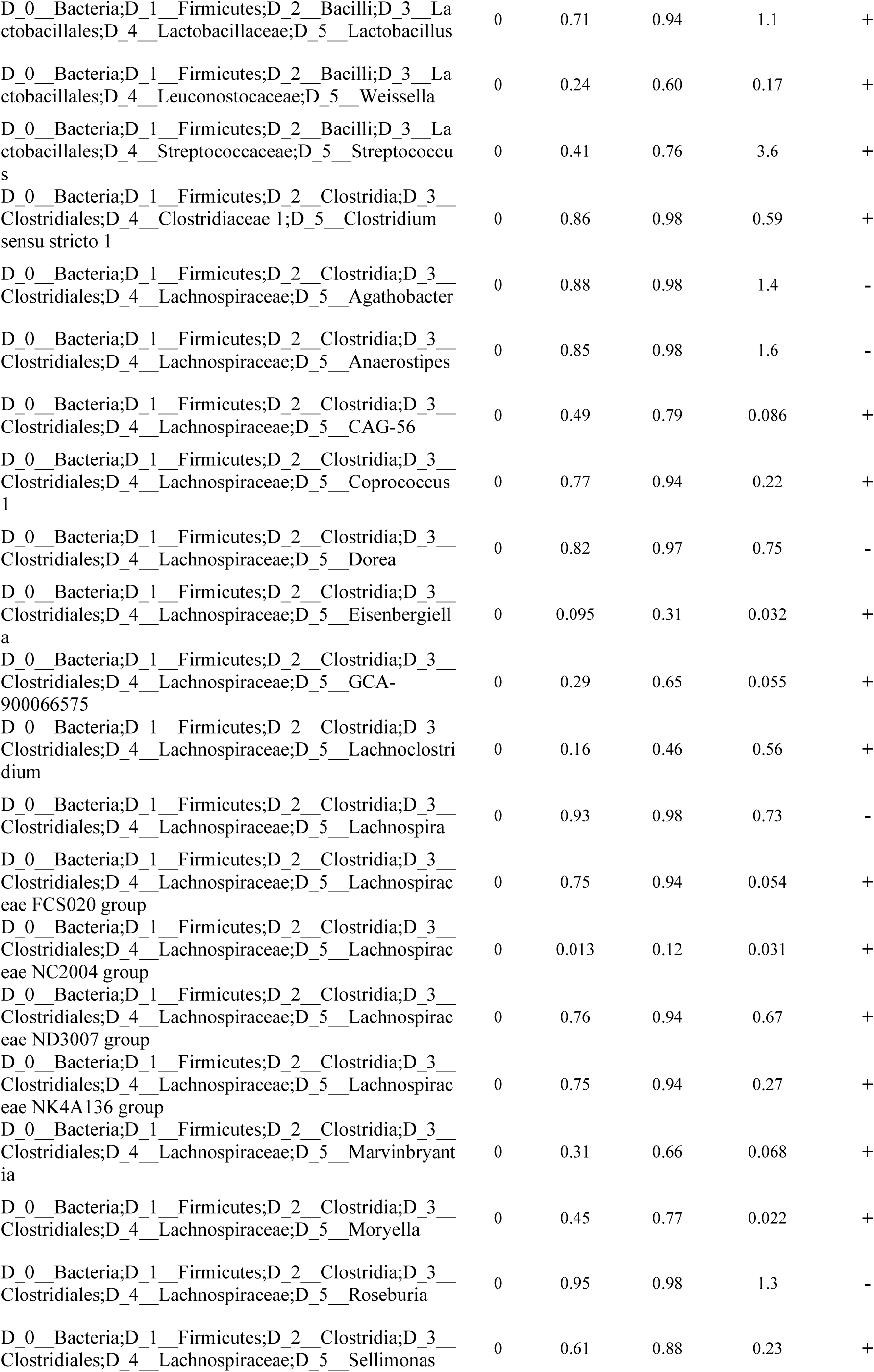
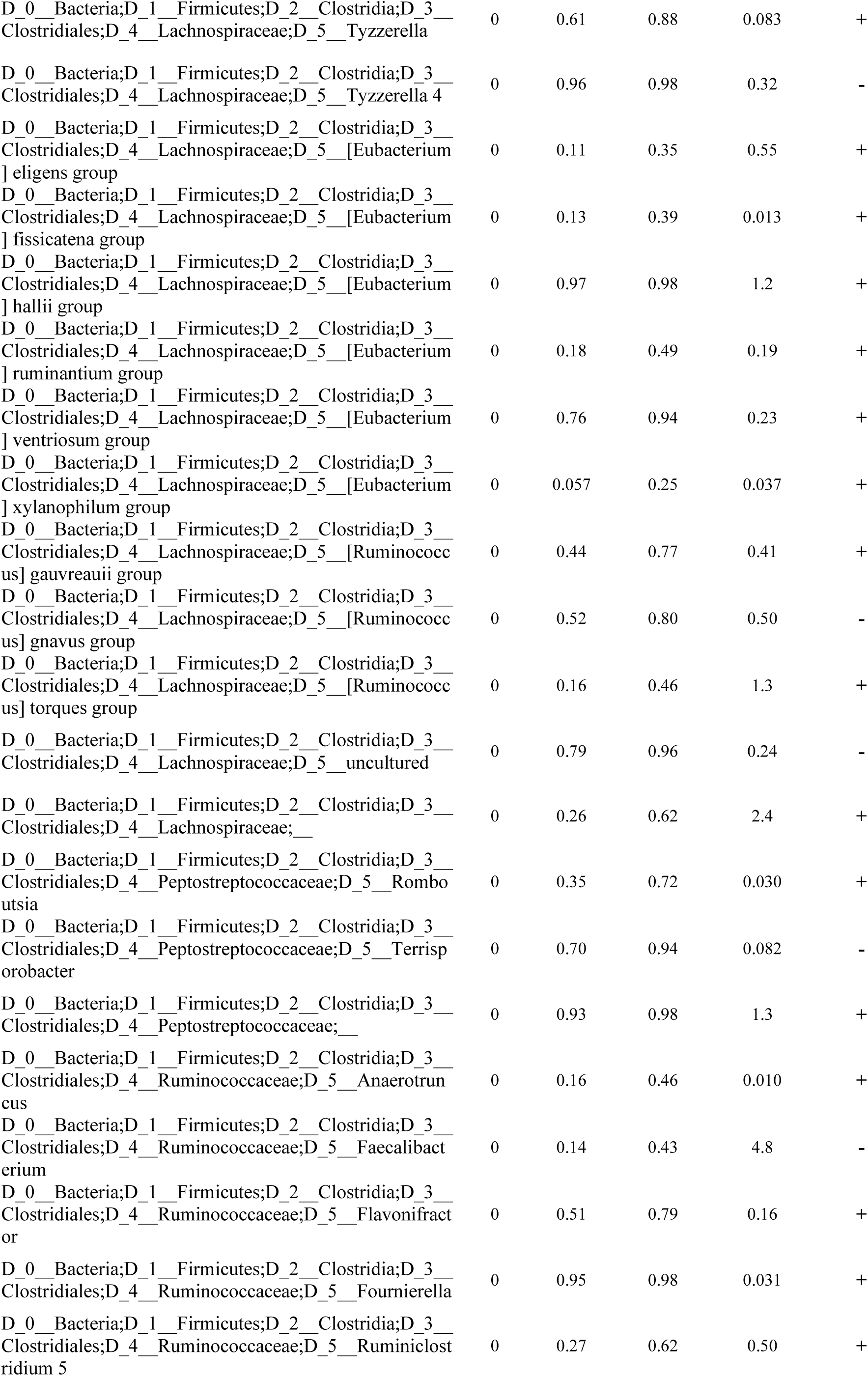
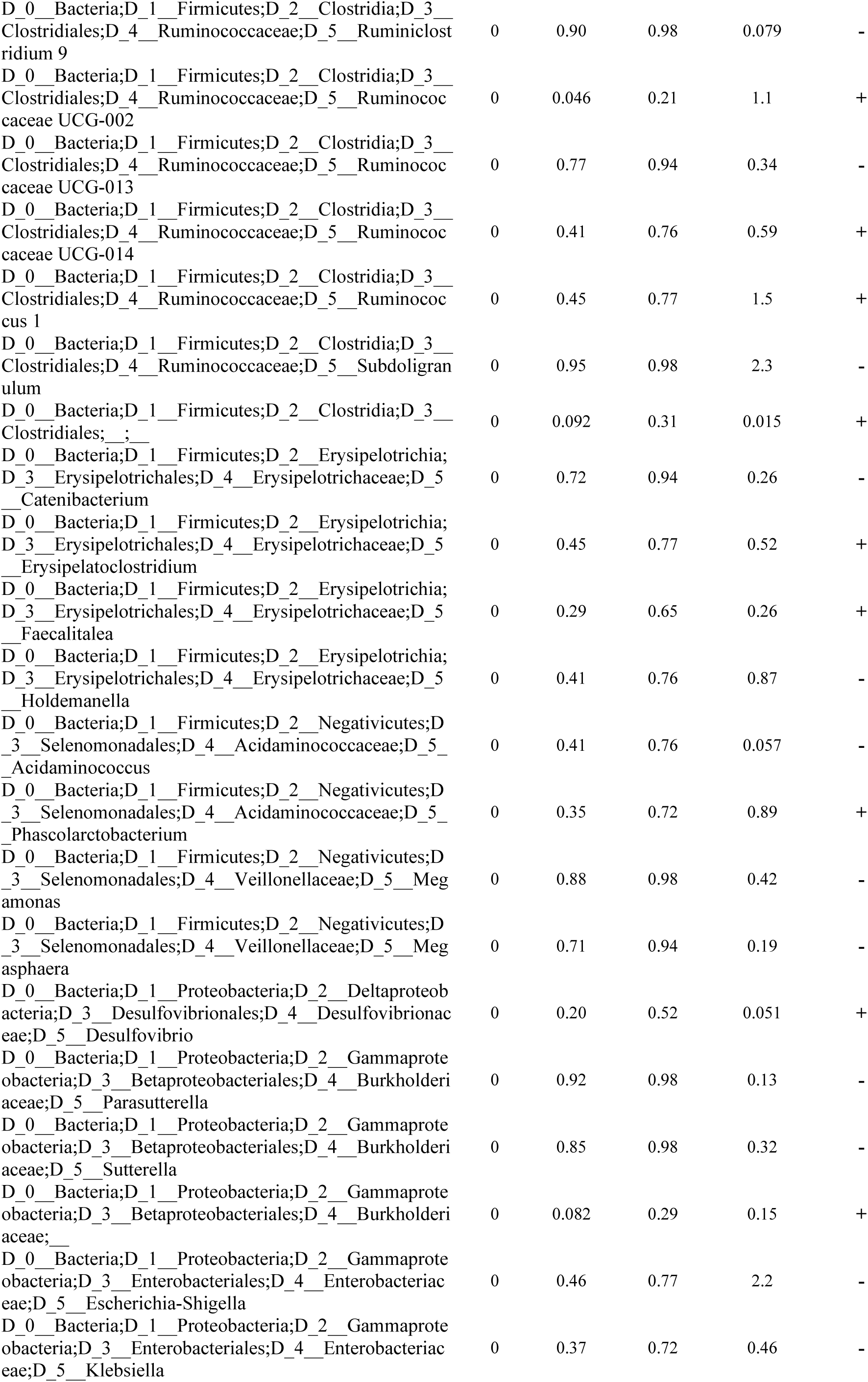
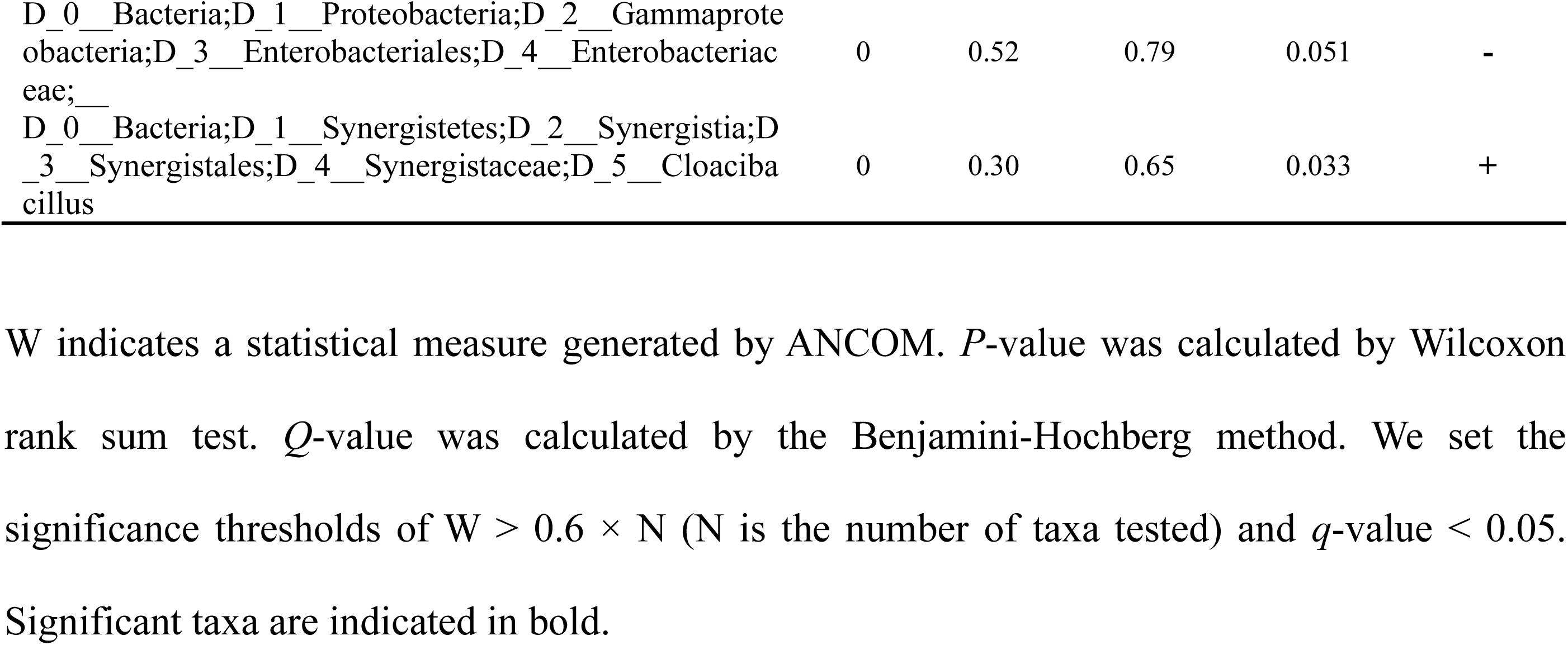
Genera changed in RBD in our dataset

**Supplementary Table S2b.**
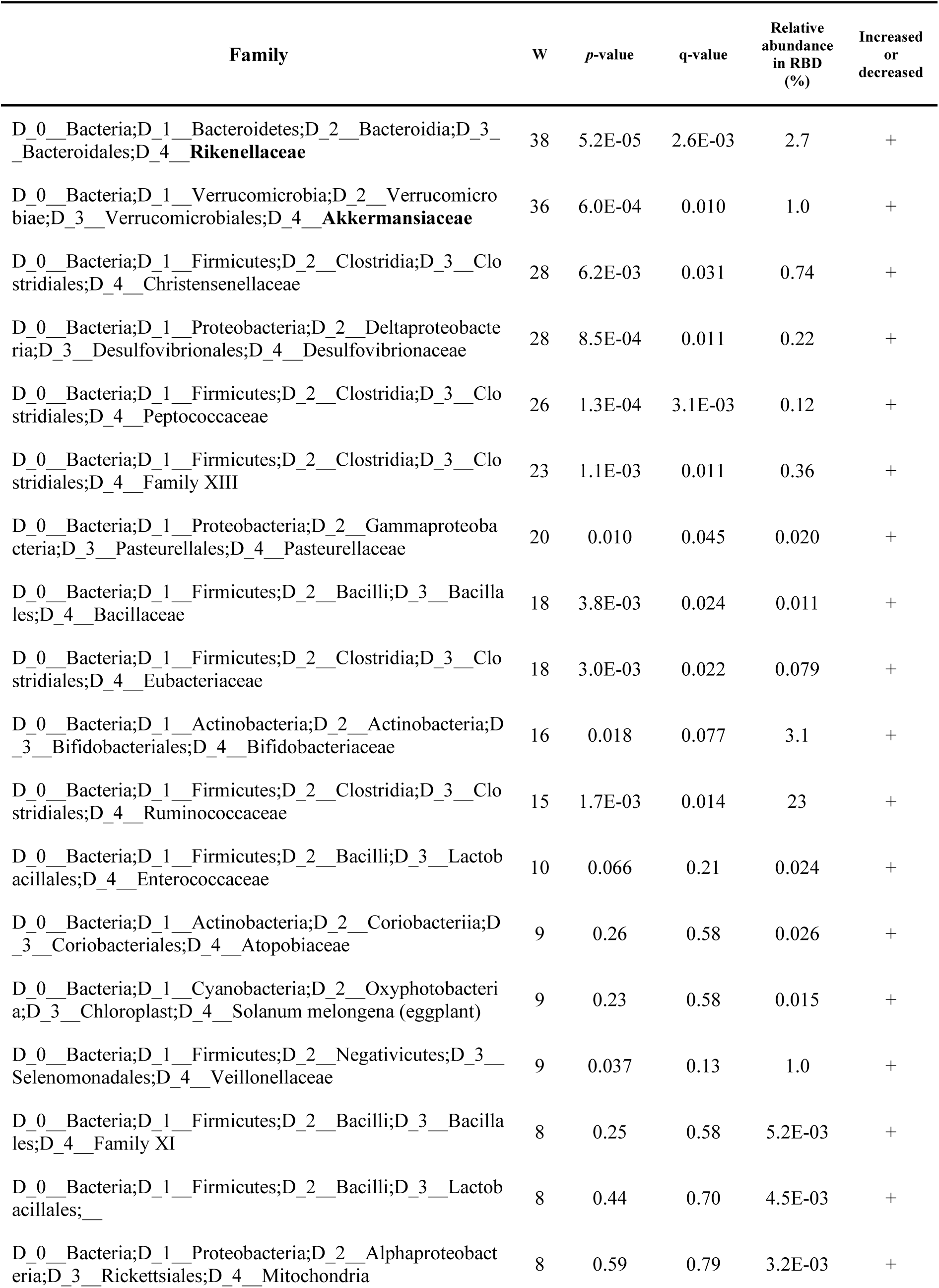
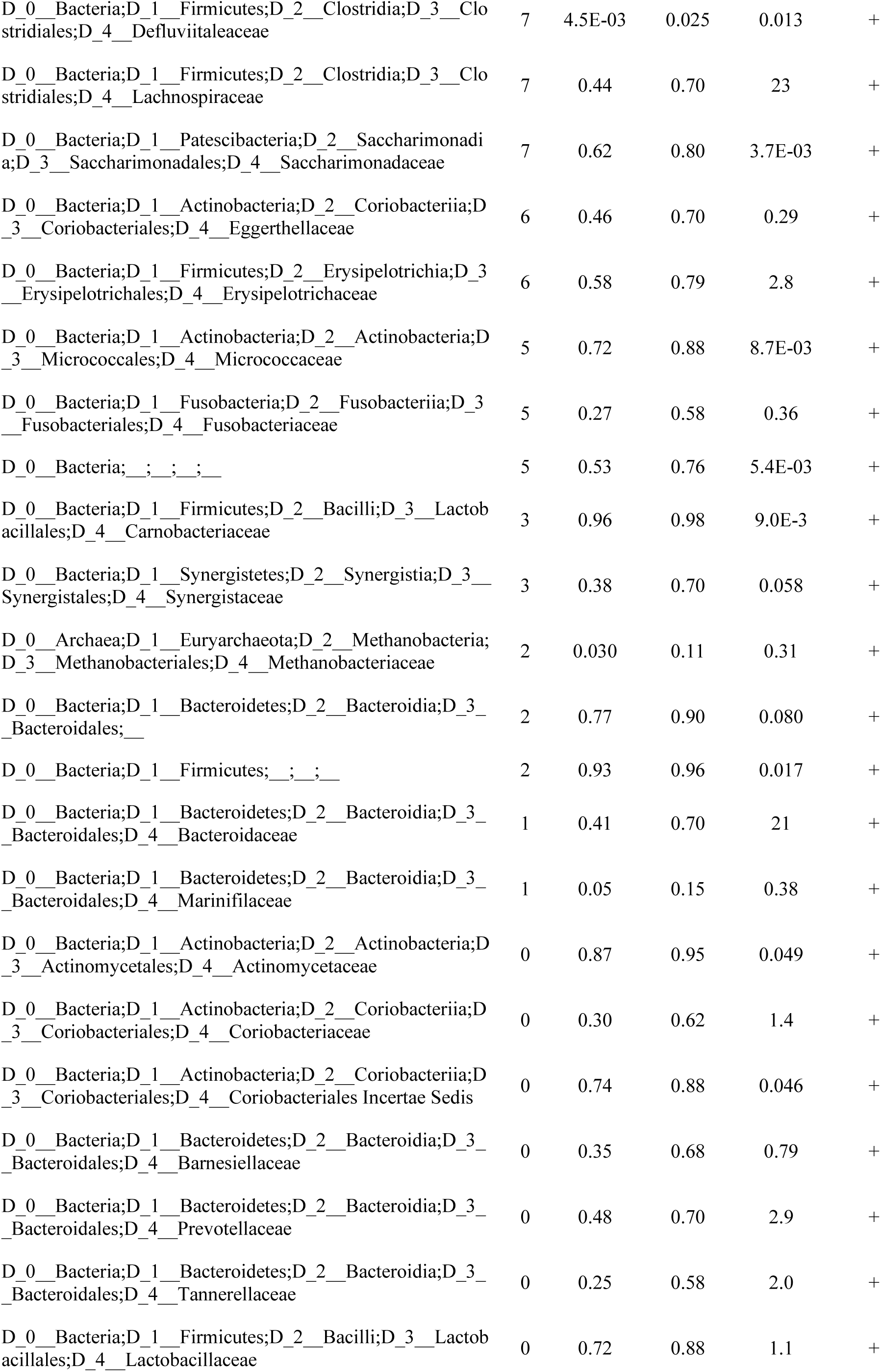
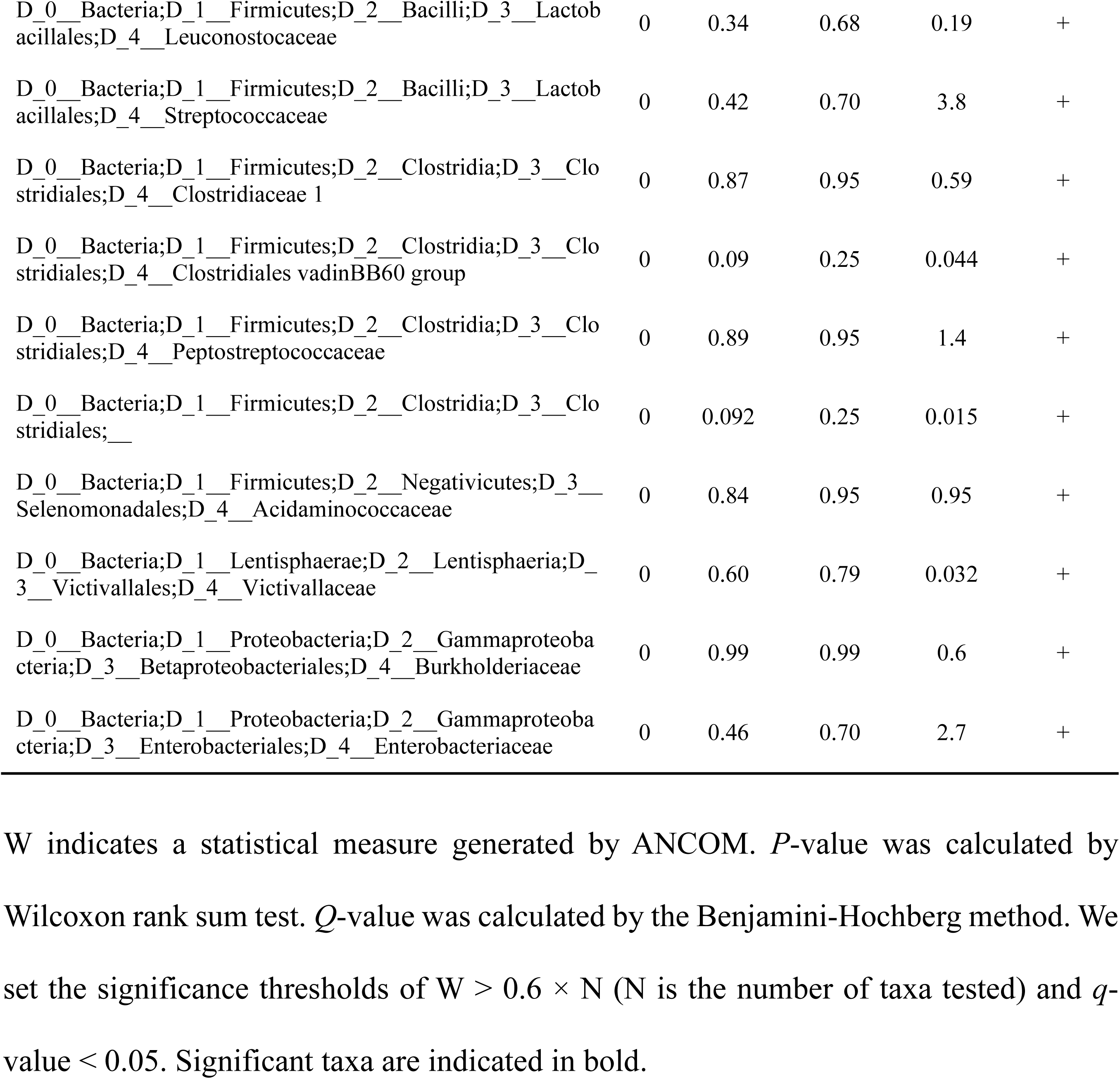
Families changed in RBD in our dataset

**Supplementary Table S3a.** Effect sizes and relative abundances of all filtered genera in RBD in the meta-analysis of the Japanese and German datasets

| Genus | Effect size | Relative abundance in RBD (%) |
| --- | --- | --- |
| D_0__Bacteria;D_1__Firmicutes;D_2__Clostridia;D_3__Clostridiales;D_4__Ruminococcaceae;D_5__Ruminococcaceae UCG-004 | 0.33 | 0.18 |
| D_0__Bacteria;D_1__Bacteroidetes;D_2__Bacteroidia;D_3__Bacteroidales;D_4__Rikenellaceae;D_5__Alistipes | 0.29 | 2.7 |
| D_0__Bacteria;D_1__Firmicutes;D_2__Clostridia;D_3__Clostridiales;D_4__Family XIII;D_5__Family XIII AD3011 group | 0.25 | 0.24 |
| D_0__Bacteria;D_1__Verrucomicrobia;D_2__Verrucomicrobiae;D_3__Verrucomicrobiales;D_4__Akkermansia;D_5__Akkermansia | 0.24 | 2.0 |
| D_0__Bacteria;D_1__Firmicutes;D_2__Clostridia;D_3__Clostridiales;D_4__Ruminococcaceae;D_5__UBA1819 | 0.24 | 0.14 |
| D_0__Bacteria;D_1__Firmicutes;D_2__Clostridia;D_3__Clostridiales;D_4__Ruminococcaceae;D_5__[Eubacterium] coprostanoligenes group | 0.23 | 1.4 |
| D_0__Bacteria;D_1__Firmicutes;D_2__Clostridia;D_3__Clostridiales;D_4__Ruminococcaceae;D_5__uncultured | 0.21 | 0.26 |
| D_0__Bacteria;D_1__Firmicutes;D_2__Clostridia;D_3__Clostridiales;D_4__Ruminococcaceae;D_5__Oscillibacter | 0.21 | 0.39 |
| D_0__Bacteria;D_1__Firmicutes;D_2__Bacilli;D_3__Bacillales;D_4__Bacillaceae;D_5__Bacillus | -0.20 | 8.6E-03 |
| D_0__Bacteria;D_1__Firmicutes;D_2__Clostridia;D_3__Clostridiales;D_4__Ruminococcaceae;D_5__Ruminococcus 2 | 0.20 | 3.8 |
| D_0__Bacteria;D_1__Firmicutes;D_2__Clostridia;D_3__Clostridiales;D_4__Lachnospiraceae;D_5__Eisenbergiella | 0.20 | 0.028 |
| D_0__Bacteria;D_1__Firmicutes;D_2__Clostridia;D_3__Clostridiales;D_4__Ruminococcaceae;D_5__Ruminococcaceae NK4A214 group | 0.19 | 0.37 |
| D_0__Bacteria;D_1__Firmicutes;D_2__Clostridia;D_3__Clostridiales;D_4__Ruminococcaceae;D_5__Anaerotruncus | 0.19 | 0.022 |
| D_0__Bacteria;D_1__Firmicutes;D_2__Clostridia;D_3__Clostridiales;D_4__Ruminococcaceae;D_5__ | 0.19 | 2.2 |
| D_0__Bacteria;D_1__Firmicutes;D_2__Clostridia;D_3__Clostridiales;D_4__Ruminococcaceae;D_5__Ruminococcaceae UCG-010 | 0.19 | 0.12 |
| D_0__Bacteria;D_1__Firmicutes;D_2__Clostridia;D_3__Clostridiales;D_4__Ruminococcaceae;D_5__Negativibacillus | 0.19 | 0.18 |
| D_0__Bacteria;D_1__Firmicutes;D_2__Erysipelotrichia;D_3__Erysipelotrichales;D_4__Erysipelotrichaceae;D_5__Holdemania | 0.18 | 0.035 |
| D_0__Bacteria;D_1__Firmicutes;D_2__Clostridia;D_3__Clostridiales;D_4__Christensenellaceae;D_5__Christensenellaceae R-7 group | 0.18 | 2.1 |
| D_0__Bacteria;D_1__Firmicutes;D_2__Clostridia;D_3__Clostridiales;D_4__Lachnospiraceae;D_5__Lachnospiraceae NC2004 group | 0.18 | 0.070 |
| D_0__Bacteria;D_1__Firmicutes;D_2__Erysipelotrichia;D_3__Erysipelotrichales;D_4__Erysipelotrichaceae;D_5__Turicibacter | -0.18 | 0.025 |
| D_0__Bacteria;D_1__Firmicutes;D_2__Clostridia;D_3__Clostridiales;D_4__Lachnospiraceae;D_5__UC5-1-2E3 | 0.17 | 0.011 |
| D_0__Bacteria;D_1__Bacteroidetes;D_2__Bacteroidia;D_3__Bacteroidales;D_4__Prevotellaceae;D_5__Prevotella 9 | -0.17 | 1.7 |
| D_0__Bacteria;D_1__Firmicutes;D_2__Clostridia;D_3__Clostridiales;D_4__Ruminococcaceae;D_5__Ruminococcaceae UCG-005 | 0.17 | 0.55 |
| D_0__Bacteria;D_1__Firmicutes;D_2__Clostridia;D_3__Clostridiales;D_4__Ruminococcaceae;D_5__Ruminiclostridium 6 | 0.17 | 0.57 |
| D_0__Bacteria;D_1__Firmicutes;D_2__Clostridia;D_3__Clostridiales;D_4__Ruminococcaceae;D_5__DTU089 | 0.16 | 0.027 |
| D_0__Bacteria;D_1__Firmicutes;D_2__Clostridia;D_3__Clostridiales;D_4__Lachnospiraceae;D_5__[Ruminococcus] torques group | 0.16 | 1.3 |
| D_0__Bacteria;D_1__Firmicutes;D_2__Clostridia;D_3__Clostridiales;D_4__Lachnospiraceae;D_5__[Eubacterium] xylanophilum group | 0.16 | 0.093 |
| D_0__Bacteria;D_1__Actinobacteria;D_2__Actinobacteria;D_3__Bifidobacteriales;D_4__Bifidobacteriaceae;D_5__Bifidobacterium | -0.16 | 2.4 |
| D_0__Bacteria;D_1__Firmicutes;D_2__Clostridia;D_3__Clostridiales;D_4__Family XIII;D_5__[Eubacterium] nodatum group | 0.14 | 0.010 |
| D_0__Bacteria;D_1__Firmicutes;D_2__Clostridia;D_3__Clostridiales;D_4__Ruminococcaceae;D_5__Faecalibacterium | -0.13 | 6.9 |
| D_0__Bacteria;D_1__Firmicutes;D_2__Clostridia;D_3__Clostridiales;D_4__Ruminococcaceae;D_5__Ruminococcaceae UCG-002 | 0.13 | 1.1 |
| D_0__Bacteria;D_1__Firmicutes;D_2__Negativicutes;D_3__Selenomonadales;D_4__Veillonellaceae;D_5__Dialister | -0.1 | 0.17 |
| D_0__Bacteria;D_1__Firmicutes;D_2__Clostridia;D_3__Clostridiales;D_4__Lachnospiraceae;D_5__Lachnoclostridium | 0.13 | 0.61 |
| D_0__Bacteria;D_1__Firmicutes;D_2__Clostridia;D_3__Clostridiales;D_4__Lachnospiraceae;D_5__Lachnospiraceae UCG-004 | -0.13 | 0.14 |
| D_0__Bacteria;D_1__Bacteroidetes;D_2__Bacteroidia;D_3__Bacteroidales;D_4__Marinifilaceae;__ | 0.13 | 0.054 |
| D_0__Bacteria;D_1__Firmicutes;D_2__Clostridia;D_3__Clostridiales;D_4__Ruminococcaceae;D_5__Ruminiclostridium 5 | 0.13 | 0.35 |
| D_0__Bacteria;D_1__Proteobacteria;D_2__Gammaproteobacteria;D_3__Betaproteobacteriales;D_4__Burkholderiaceae;__ | 0.12 | 0.62 |
| D_0__Bacteria;D_1__Firmicutes;D_2__Clostridia;D_3__Clostridiales;D_4__Lachnospiraceae;__ | 0.12 | 4.1 |
| D_0__Bacteria;D_1__Bacteroidetes;D_2__Bacteroidia;D_3__Bacteroidales;D_4__Tannerellaceae;D_5__Parabacteroides | 0.11 | 2.1 |
| D_0__Bacteria;D_1__Firmicutes;D_2__Clostridia;D_3__Clostridiales;D_4__Lachnospiraceae;D_5__Coproccoccus 2 | -0.11 | 0.24 |
| D_0__Bacteria;D_1__Firmicutes;D_2__Clostridia;D_3__Clostridiales;__;__ | 0.11 | 0.27 |
| D_0__Bacteria;D_1__Firmicutes;D_2__Clostridia;D_3__Clostridiales;D_4__Lachnospiraceae;D_5__Marvinbryantia | 0.11 | 0.10 |
| D_0__Archaea;D_1__Euryarchaeota;D_2__Methanobacteria;D_3__Methanobacteriales;D_4__Methanobacteriaceae;D_5__Methanobrevibacter | 0.11 | 0.21 |
| D_0__Bacteria;D_1__Bacteroidetes;D_2__Bacteroidia;D_3__Bacteroidales;D_4__Marinifilaceae;D_5__Butyricimonas | 0.11 | 0.14 |
| D_0__Bacteria;D_1__Firmicutes;D_2__Clostridia;D_3__Clostridiales;D_4__Ruminococcaceae;D_5__Butyricicoccus | -0.10 | 0.29 |
| D_0__Bacteria;D_1__Bacteroidetes;D_2__Bacteroidia;D_3__Bacteroidales;D_4__Barnesiellaceae;D_5__Barnesiella | 0.10 | 0.68 |
| D_0__Bacteria;D_1__Firmicutes;D_2__Erysipelotrichia;D_3__Erysipelotrichales;D_4__Erysipelotrichaceae;D_5__Erysipelatoclostridium | 0.10 | 0.33 |
| D_0__Bacteria;D_1__Firmicutes;D_2__Clostridia;D_3__Clostridiales;D_4__Ruminococcaceae;D_5__Flavonifractor | 0.10 | 0.16 |
| D_0__Bacteria;D_1__Firmicutes;D_2__Clostridia;D_3__Clostridiales;D_4__Ruminococcaceae;D_5__Intestinimonas | 0.091 | 0.079 |
| D_0__Bacteria;D_1__Bacteroidetes;D_2__Bacteroidia;D_3__Bacteroidales;D_4__Marinifilaceae;D_5__Odoribacter | 0.090 | 0.31 |
| D_0__Bacteria;D_1__Firmicutes;D_2__Erysipelotrichia;<br>D_3__Erysipelotrichales;D_4__Erysipelotrichaceae;D_5__<br>uncultured | 0.089 | 0.027 |
| D_0__Bacteria;D_1__Firmicutes;D_2__Erysipelotrichia;<br>D_3__Erysipelotrichales;D_4__Erysipelotrichaceae;__ | 0.089 | 0.037 |
| D_0__Bacteria;D_1__Actinobacteria;D_2__Coriobacterii<br>a;D_3__Coriobacteriales;D_4__Eggerthellaceae;D_5__Sl<br>ackia | 0.085 | 0.053 |
| D_0__Bacteria;D_1__Firmicutes;D_2__Clostridia;D_3__<br>Clostridiales;D_4__Ruminococcaceae;D_5__Ruminococc<br>aceae UCG-014 | 0.081 | 0.48 |
| D_0__Bacteria;D_1__Actinobacteria;D_2__Coriobacterii<br>a;D_3__Coriobacteriales;D_4__Eggerthellaceae;__ | 0.079 | 0.067 |
| D_0__Bacteria;D_1__Firmicutes;D_2__Clostridia;D_3__<br>Clostridiales;D_4__Lachnospiraceae;D_5__Sellimonas | 0.078 | 0.14 |
| D_0__Bacteria;D_1__Bacteroidetes;D_2__Bacteroidia;D<br>_3__Bacteroidales;D_4__Prevotellaceae;D_5__Prevotella<br>2 | 0.077 | 0.37 |
| D_0__Bacteria;D_1__Bacteroidetes;D_2__Bacteroidia;D<br>_3__Bacteroidales;D_4__Barnesiellaceae;D_5__unculture<br>d | 0.077 | 0.083 |
| D_0__Bacteria;D_1__Patescibacteria;D_2__Saccharimon<br>adia;D_3__Saccharimonadales;D_4__Saccharimonadacea<br>e;D_5__TM7 phylum sp. canine oral taxon 250 | -0.073 | 2.5E-03 |
| D_0__Bacteria;D_1__Firmicutes;D_2__Clostridia;D_3__<br>Clostridiales;D_4__Lachnospiraceae;D_5__[Eubacterium]<br>eligans group | 0.072 | 0.53 |
| D_0__Bacteria;D_1__Firmicutes;D_2__Clostridia;D_3__<br>Clostridiales;D_4__Peptostreptococcaceae;D_5__Rombou<br>tsia | 0.072 | 0.018 |
| D_0__Bacteria;D_1__Firmicutes;D_2__Clostridia;D_3__<br>Clostridiales;D_4__Lachnospiraceae;D_5__Coproccoccus<br>1 | 0.072 | 0.23 |
| D_0__Bacteria;D_1__Bacteroidetes;D_2__Bacteroidia;D<br>_3__Bacteroidales;D_4__Prevotellaceae;D_5__Paraprevo<br>tella | 0.071 | 0.27 |
| D_0__Bacteria;D_1__Firmicutes;D_2__Negativicutes;D_<br>3__Selenomonadales;D_4__Veillonellaceae;D_5__Megas<br>phaera | -0.071 | 0.11 |
| D_0__Bacteria;D_1__Firmicutes;D_2__Clostridia;D_3__<br>Clostridiales;D_4__Lachnospiraceae;D_5__Lachnospirac<br>eae UCG-008 | -0.066 | 0.042 |
| D_0__Bacteria;D_1__Firmicutes;D_2__Clostridia;D_3__<br>Clostridiales;D_4__Lachnospiraceae;D_5__Blautia | 0.066 | 4.7 |
| D_0__Bacteria;D_1__Actinobacteria;D_2__Coriobacterii<br>a;D_3__Coriobacteriales;D_4__Eggerthellaceae;D_5__un<br>cultured | -0.066 | 0.043 |
| D_0__Bacteria;D_1__Firmicutes;D_2__Clostridia;D_3__<br>Clostridiales;D_4__Ruminococcaceae;D_5__Candidatus<br>Soleaferrea | -0.064 | 4.6E-03 |
| D_0__Bacteria;D_1__Firmicutes;D_2__Erysipelotrichia;<br>D_3__Erysipelotrichales;D_4__Erysipelotrichaceae;D_5__<br>_Catenibacterium | -0.064 | 0.28 |
| D_0__Bacteria;D_1__Firmicutes;D_2__Erysipelotrichia;<br>D_3__Erysipelotrichales;D_4__Erysipelotrichaceae;D_5__<br>_Merdibacter | -0.063 | 6.7E-03 |
| D_0__Bacteria;D_1__Firmicutes;D_2__Clostridia;D_3__<br>Clostridiales;D_4__Ruminococcaceae;D_5__Ruminococc<br>aceae UCG-003 | -0.062 | 0.12 |
| D_0__Bacteria;D_1__Firmicutes;D_2__Clostridia;D_3__<br>Clostridiales;D_4__Clostridiaceae 1;D_5__Clostridium<br>sensu stricto 1 | -0.062 | 0.40 |
| D_0__Bacteria;D_1__Firmicutes;D_2__Bacilli;D_3__Lac<br>tobacillales;D_4__Streptococcaceae;D_5__Streptococcus | 0.059 | 2.2 |
| D_0__Bacteria;D_1__Bacteroidetes;D_2__Bacteroidia;D<br>_3__Bacteroidales;D_4__Barnesiellaceae;D_5__Coproba<br>cter | 0.059 | 0.087 |
| D_0__Bacteria;D_1__Firmicutes;D_2__Clostridia;D_3__<br>Clostridiales;D_4__Lachnospiraceae;D_5__Lachnospira | -0.058 | 0.66 |
| D_0__Bacteria;D_1__Synergistetes;D_2__Synergistia;D_<br>3__Synergistales;D_4__Synergistaceae;D_5__Cloacibacil<br>lus | 0.056 | 0.034 |
| D_0__Bacteria;D_1__Firmicutes;D_2__Clostridia;D_3__<br>Clostridiales;D_4__Lachnospiraceae;D_5__Fusicatenibact<br>er | -0.056 | 1.1 |
| D_0__Bacteria;D_1__Firmicutes;D_2__Clostridia;D_3__<br>Clostridiales;D_4__Lachnospiraceae;D_5__Lachnospirac<br>eae ND3007 group | 0.055 | 0.76 |
| D_0__Bacteria;D_1__Firmicutes;D_2__Clostridia;D_3__<br>Clostridiales;D_4__Ruminococcaceae;D_5__GCA-900066225 | 0.054 | 0.013 |
| D_0__Bacteria;D_1__Actinobacteria;D_2__Coriobacterii<br>a;D_3__Coriobacteriales;D_4__Coriobacteriaceae;D_5__<br>Collinsella | -0.051 | 1.1 |
| D_0__Bacteria;D_1__Firmicutes;D_2__Clostridia;D_3__<br>Clostridiales;D_4__Lachnospiraceae;D_5__Tyzzerella | 0.048 | 0.061 |
| D_0__Bacteria;D_1__Firmicutes;D_2__Clostridia;D_3__<br>Clostridiales;D_4__Lachnospiraceae;D_5__Lachnospirac<br>eae UCG-010 | -0.048 | 0.039 |
| D_0__Bacteria;D_1__Firmicutes;D_2__Erysipelotrichia;<br>D_3__Erysipelotrichales;D_4__Erysipelotrichaceae;D_5__<br>_Erysipelotrichaceae UCG-003 | -0.048 | 0.48 |
| D_0__Bacteria;D_1__Actinobacteria;D_2__Coriobacterii<br>a;D_3__Coriobacteriales;D_4__Eggerthellaceae;D_5__Eg<br>gerthella | -0.048 | 0.091 |
| D_0__Bacteria;D_1__Actinobacteria;D_2__Coriobacterii<br>a;D_3__Coriobacteriales;D_4__Coriobacteriales Incertae<br>Sedis;D_5__uncultured | 0.047 | 0.040 |
| D_0__Bacteria;D_1__Firmicutes;D_2__Clostridia;D_3__Clostridiales;D_4__Lachnospiraceae;D_5__[Ruminococcus] gnavus group | 0.047 | 0.35 |
| D_0__Bacteria;D_1__Firmicutes;D_2__Erysipelotrichia;D_3__Erysipelotrichales;D_4__Erysipelotrichaceae;D_5__[Clostridium] innocuum group | 0.047 | 0.049 |
| D_0__Bacteria;D_1__Firmicutes;D_2__Clostridia;D_3__Clostridiales;D_4__Lachnospiraceae;D_5__[Eubacterium] ruminantium group | 0.046 | 0.38 |
| D_0__Bacteria;D_1__Firmicutes;D_2__Erysipelotrichia;D_3__Erysipelotrichales;D_4__Erysipelotrichaceae;D_5__Faecalitalea | 0.046 | 0.17 |
| D_0__Bacteria;D_1__Firmicutes;D_2__Clostridia;D_3__Clostridiales;D_4__Lachnospiraceae;D_5__Dorea | -0.043 | 0.63 |
| D_0__Bacteria;D_1__Firmicutes;D_2__Clostridia;D_3__Clostridiales;D_4__Ruminococcaceae;D_5__Phoceia | 0.043 | 5.0E-03 |
| D_0__Bacteria;D_1__Proteobacteria;D_2__Gammaproteobacteria;D_3__Enterobacteriales;D_4__Enterobacteriaceae;D_5__Klebsiella | -0.043 | 0.28 |
| D_0__Bacteria;D_1__Firmicutes;D_2__Clostridia;D_3__Clostridiales;D_4__Lachnospiraceae;D_5__[Ruminococcus] gauvreauii group | 0.043 | 0.38 |
| D_0__Bacteria;D_1__Firmicutes;D_2__Clostridia;D_3__Clostridiales;D_4__Peptostreptococcaceae;D_5__ | -0.043 | 0.75 |
| D_0__Bacteria;D_1__Proteobacteria;D_2__Deltaproteobacteria;D_3__Desulfovibrionales;D_4__Desulfovibrionaceae;D_5__Desulfovibrio | 0.040 | 0.29 |
| D_0__Bacteria;D_1__Firmicutes;D_2__Clostridia;D_3__Clostridiales;D_4__Ruminococcaceae;D_5__Ruminococcaceae UCG-009 | 0.039 | 1.1E-02 |
| D_0__Bacteria;D_1__Firmicutes;D_2__Negativicutes;D_3__Selenomonadales;D_4__Acidaminococcaceae;D_5__Phascolarctobacterium | 0.032 | 0.86 |
| D_0__Bacteria;D_1__Firmicutes;D_2__Clostridia;D_3__Clostridiales;D_4__Ruminococcaceae;D_5__Ruminococcaceae UCG-013 | 0.032 | 0.24 |
| D_0__Bacteria;D_1__Firmicutes;D_2__Clostridia;D_3__Clostridiales;D_4__Lachnospiraceae;D_5__Agathobacter | -0.032 | 1.7 |
| D_0__Bacteria;D_1__Firmicutes;D_2__Clostridia;D_3__Clostridiales;D_4__Lachnospiraceae;D_5__Roseburia | 0.032 | 1.4 |
| D_0__Bacteria;D_1__Firmicutes;D_2__Clostridia;D_3__Clostridiales;D_4__Lachnospiraceae;D_5__[Eubacterium] hallii group | 0.031 | 1.2 |
| D_0__Bacteria;D_1__Firmicutes;D_2__Erysipelotrichia;D_3__Erysipelotrichales;D_4__Erysipelotrichaceae;D_5__Coprobacillus | 0.030 | 0.019 |
| D_0__Bacteria;D_1__Bacteroidetes;D_2__Bacteroidia;D_3__Bacteroidales;D_4__Prevotellaceae;D_5__ | -0.028 | 0.27 |
| D_0__Bacteria;D_1__Firmicutes;D_2__Bacilli;D_3__Lactobacillales;D_4__Streptococcaceae;D_5__Lactococcus | 0.027 | 0.12 |
| D_0__Bacteria;D_1__Fusobacteria;D_2__Fusobacteriia;D_3__Fusobacteriales;D_4__Fusobacteriaceae;D_5__Fusobacterium | -0.027 | 0.20 |
| D_0__Bacteria;D_1__Firmicutes;D_2__Clostridia;D_3__Clostridiales;D_4__Lachnospiraceae;D_5__[Eubacterium] ventriosum group | 0.026 | 0.14 |
| D_0__Bacteria;D_1__Firmicutes;D_2__Clostridia;D_3__Clostridiales;D_4__Ruminococcaceae;D_5__Ruminiclostridium 9 | -0.024 | 0.11 |
| D_0__Bacteria;D_1__Bacteroidetes;D_2__Bacteroidia;D_3__Bacteroidales;D_4__Prevotellaceae;D_5__Alloprevotella | -0.024 | 0.12 |
| D_0__Bacteria;D_1__Actinobacteria;D_2__Actinobacteriia;D_3__Actinomycetales;D_4__Actinomycetaceae;D_5__Actinomyces | 0.023 | 0.033 |
| D_0__Bacteria;D_1__Actinobacteria;D_2__Coriobacteriia;D_3__Coriobacteriales;D_4__Eggerthellaceae;D_5__Gordonibacter | 0.023 | 0.012 |
| D_0__Bacteria;__;__;__;__; | -0.023 | 0.076 |
| D_0__Bacteria;D_1__Bacteroidetes;D_2__Bacteroidia;D_3__Bacteroidales;D_4__Prevotellaceae;D_5__uncultured | 0.023 | 0.073 |
| D_0__Bacteria;D_1__Firmicutes;D_2__Bacilli;D_3__Lactobacillales;D_4__Lactobacillaceae;D_5__Lactobacillus | 0.021 | 0.91 |
| D_0__Bacteria;D_1__Firmicutes;D_2__Clostridia;D_3__Clostridiales;D_4__Lachnospiraceae;D_5__Moryella | 0.021 | 0.019 |
| D_0__Bacteria;D_1__Firmicutes;D_2__Clostridia;D_3__Clostridiales;D_4__Lachnospiraceae;D_5__Tyzzerella 3 | -0.019 | 0.040 |
| D_0__Bacteria;D_1__Bacteroidetes;D_2__Bacteroidia;D_3__Bacteroidales;__;__ | 0.018 | 0.42 |
| D_0__Bacteria;D_1__Firmicutes;D_2__Clostridia;D_3__Clostridiales;D_4__Peptostreptococcaceae;D_5__Terrisporobacter | -0.017 | 0.062 |
| D_0__Bacteria;D_1__Firmicutes;D_2__Clostridia;D_3__Clostridiales;D_4__Ruminococcaceae;D_5__Ruminococcus 1 | -0.016 | 1.2 |
| D_0__Bacteria;D_1__Firmicutes;D_2__Clostridia;D_3__Clostridiales;D_4__Lachnospiraceae;D_5__Lachnospiraceae FCS020 group | 0.012 | 0.077 |
| D_0__Bacteria;D_1__Firmicutes;__;__;__;__ | 0.012 | 0.26 |
| D_0__Bacteria;D_1__Actinobacteria;D_2__Coriobacteriia;D_3__Coriobacteriales;D_4__Eggerthellaceae;D_5__Selenigalimassilia | -7.7E-03 | 0.05 |
| D_0__Bacteria;D_1__Firmicutes;D_2__Clostridia;D_3__Clostridiales;D_4__Lachnospiraceae;D_5__CAG-56 | 7.0E-03 | 0.11 |
| D_0__Bacteria;D_1__Firmicutes;D_2__Erysipelotrichia;<br>D_3__Erysipelotrichales;D_4__Erysipelotrichaceae;D_5__<br>_Holdemanella | 6.8E-03 | 0.75 |
| D_0__Bacteria;D_1__Firmicutes;D_2__Clostridia;D_3__<br>Clostridiales;D_4__Lachnospiraceae;D_5__Lachnospirac<br>eae NK4A136 group | -6.3E-03 | 0.47 |
| D_0__Bacteria;D_1__Firmicutes;D_2__Clostridia;D_3__<br>Clostridiales;D_4__Ruminococcaceae;D_5__Oscillospira | 5.5E-03 | 0.045 |
| D_0__Bacteria;D_1__Firmicutes;D_2__Clostridia;D_3__<br>Clostridiales;D_4__Lachnospiraceae;D_5__Anaerostipes | -4.7E-03 | 1.3 |
| D_0__Bacteria;D_1__Actinobacteria;D_2__Coriobacterii<br>a;D_3__Coriobacteriales;D_4__Atopobiaceae;D_5__Olse<br>nella | -4.6E-03 | 0.023 |
| D_0__Bacteria;D_1__Bacteroidetes;D_2__Bacteroidia;D<br>_3__Bacteroidales;D_4__Bacteroidaceae;D_5__Bacteroid<br>es | 4.2E-03 | 19 |
| D_0__Bacteria;D_1__Firmicutes;D_2__Clostridia;D_3__<br>Clostridiales;D_4__Lachnospiraceae;D_5__uncultured | -4.1E-03 | 0.17 |
| D_0__Bacteria;D_1__Proteobacteria;D_2__Gammaproteo<br>bacteria;D_3__Enterobacteriales;D_4__Enterobacteriacea<br>e;__ | 7.2E-04 | 0.03 |
| D_0__Bacteria;D_1__Firmicutes;D_2__Clostridia;D_3__<br>Clostridiales;D_4__Ruminococcaceae;D_5__Subdoligran<br>ulum | 7.1E-04 | 2.2 |
| D_0__Bacteria;D_1__Firmicutes;D_2__Clostridia;D_3__<br>Clostridiales;D_4__Lachnospiraceae;D_5__GCA-900066575 | 9.5E-05 | 0.059 |

**Supplementary Table S3b.**
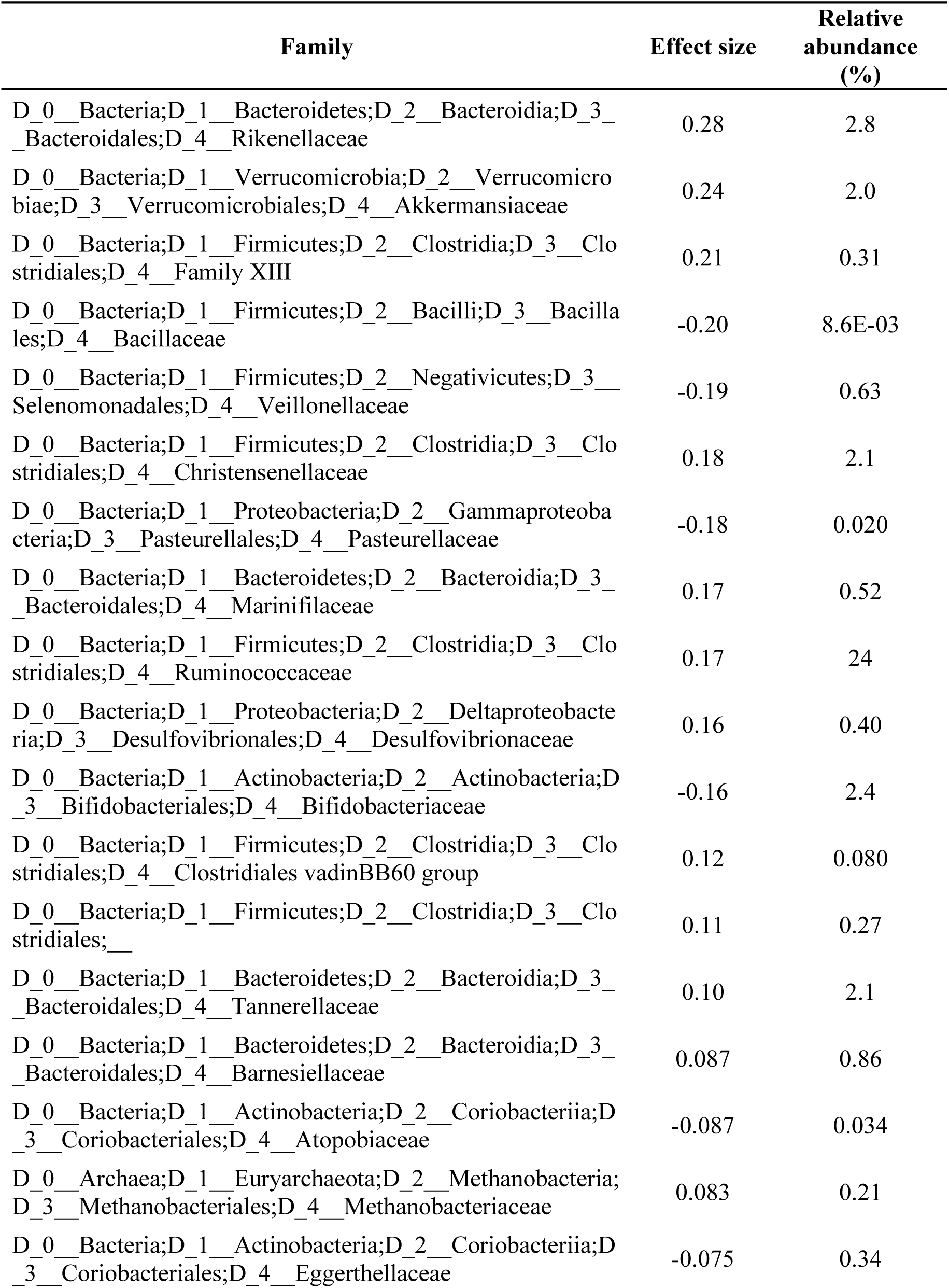
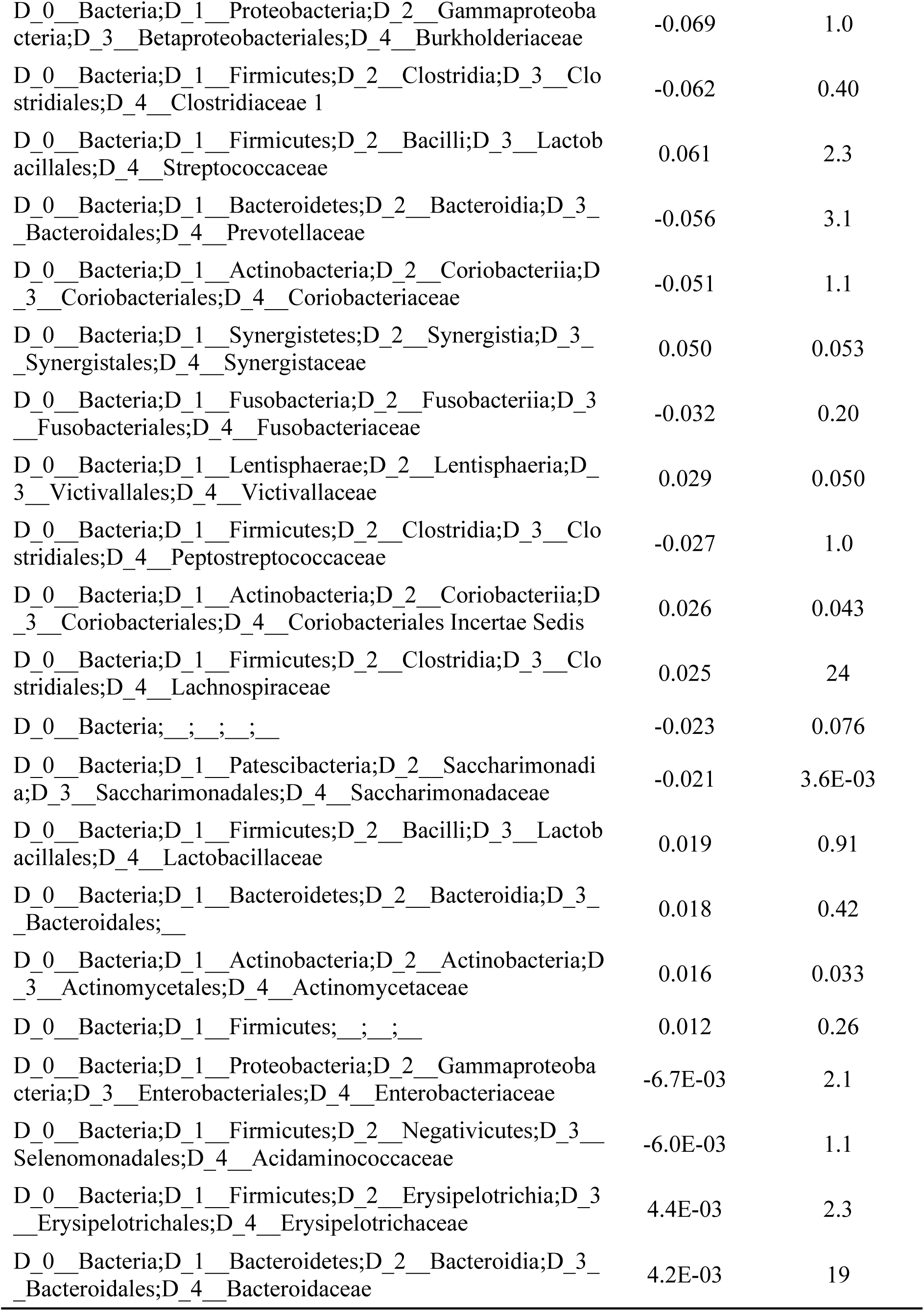
Effect sizes and relative abundances of all filtered families in RBD in the meta-analysis of the Japanese and German datasets

**Supplementary Table S4.** Experimental methods and disease durations of the Japanese and German datasets

| **Country** | **Transportation temperature** | **Storage method** | **Stool DNA stabilizer** | **Sequencing** | **Primers** | **Disease duration (years)<sup>a</sup>** |
| --- | --- | --- | --- | --- | --- | --- |
| Japan | 0 °C | Freeze dry | no | 16S rRNA V3-4 | 341F/805R | 6.4 ± 4.8<br>(Max 20, Min 0.1) |
| Germany | On dry ice | −80 °C | no | 16S rRNA V4 | 515F/805R | n.a. |

<sup>a</sup>Mean and SD. n.a., not available.

**Supplementary Table S5.** The numbers of read counts in the Japanese and German datasets

| (x 1000 read counts) |  |  |  |  |  |
| --- | --- | --- | --- | --- | --- |
| Country (sample size) | Average | SD | Median | Max | Min |
| Japan (163) | 53.5 | 18.7 | 50.8 | 98.0 | 13.8 |
| Germany (58) | 155.7 | 39.5 | 164.8 | 261.4 | 63.7 |

**Supplementary Figure S1.**
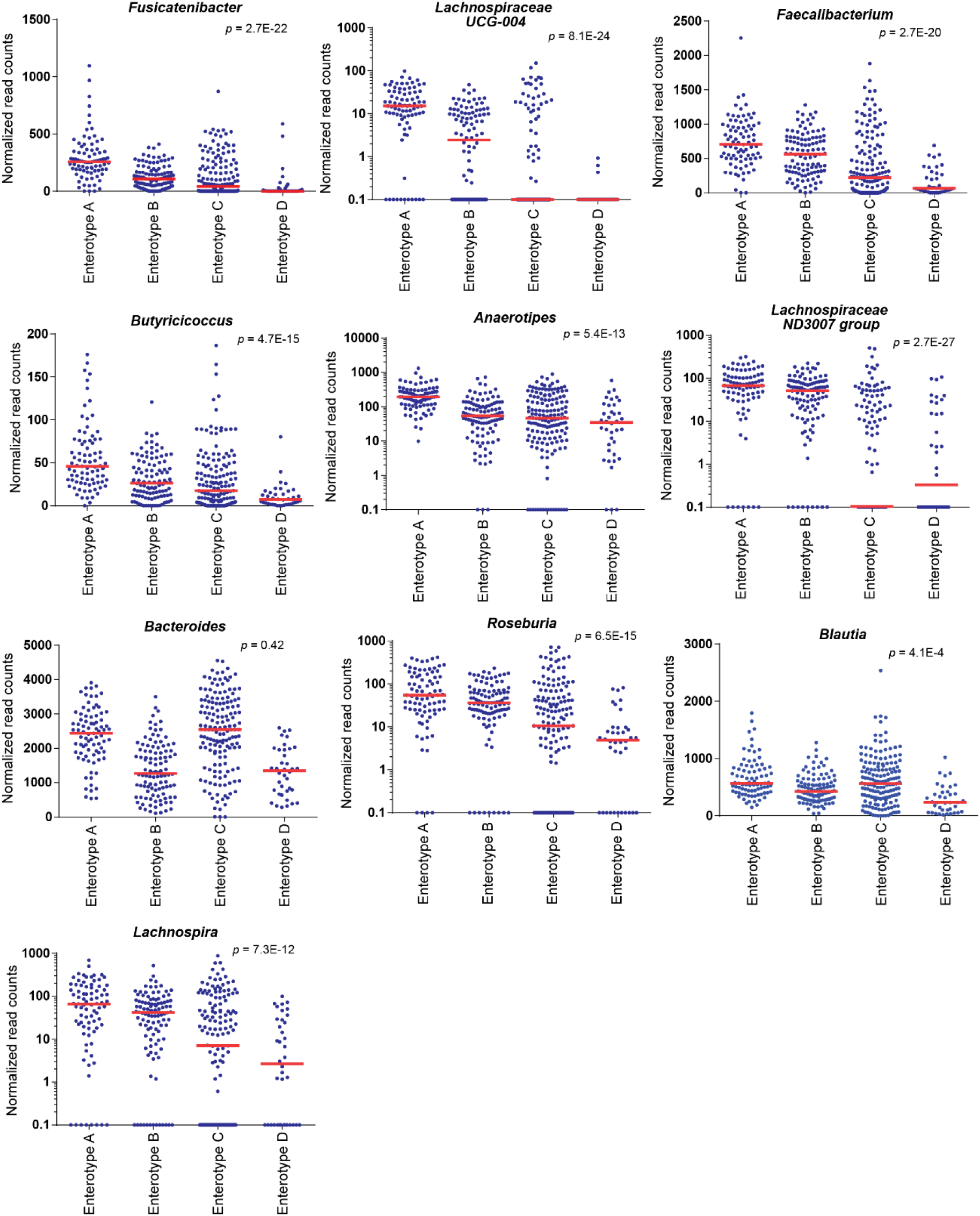
Relative abundances of ten genera with the highest loadings in the first factor (Supplementary Table S3) are plotted against enterotypes A to D generated by LIGER. Bar indicates the median value. *P*-values of Jonckheere-Terpstra trend test are indicated to show whether the genus increases or decreases monotonically.

